# Bypassing cisplatin resistance in Nrf2 hyperactivated head and neck cancer through effective PI3Kinase targeting

**DOI:** 10.1101/2025.01.10.632413

**Authors:** P. Yadollahi, K.A. McCord, Y. Li, H. Dayoub, K. Saab, F. Essien, S. Hyslop, E. Kan, K.M. Ahmed, P. R. Kirby, V. Putluri, C.S.R Ambati, K.R. Kami Reddy, P. Castro, H.D. Skinner, C. Coarfa, W.K. Decker, A.A Osman, R. Patel, J.N. Myers, S.Y. Lai, N. Putluri, F.M. Johnson, M.J. Frederick, W.H. Hudson, V.C. Sandulache

**Affiliations:** Bobby R. Alford Department of Otolaryngology Head and Neck Surgery, Baylor College of Medicine, Houston, TX; Department of Molecular and Cellular Biology, Baylor College of Medicine, Houston, TX; Department of Radiation Oncology, Baylor College of Medicine, Houston, TX; Dan L Duncan Comprehensive Cancer Center, Baylor College of Medicine, Houston, TX; Advanced Technology Cores, Baylor College of Medicine, Houston, TX; Department of Pathology and Immunology, Baylor College of Medicine, Houston, TX; Department of Radiation Oncology, UPMC Hillman Cancer Center, Pittsburgh, PA; Department of Head and Neck Surgery, UT MD Anderson Cancer Center, Houston, TX; Department of Thoracic-Head and Neck Medical Oncology, UT MD Anderson Cancer Center, Houston, TX; Center for Cell and Gene Therapy, Baylor College of Medicine, Houston, TX; Center for Translational Research on Inflammatory Diseases, Michael E. DeBakey Veterans Affairs Medical Center, Houston, TX

**Keywords:** NRF2, gedatolisib, copanlisib, head and neck cancer

## Abstract

**Background:** For patients with head and neck squamous cell carcinoma (HNSCC), failure of definitive radiation combined with cisplatin nearly universally results in death. Although hyperactivation of the Nrf2 pathway can drive radiation and cisplatin resistance along with suppressed anti-tumor immunity, treatment-refractory HNSCC tumors may retain sensitivity to targeted agents secondary to synergistic lethality with other oncogenic drivers (e.g., *NOTCH1* mutations).

**Purpose:** We evaluated the efficacy of PI3K inhibitors (PI3Ki) in bypassing Nrf2-mediated cisplatin resistance in HNSCC.

**Methods:** We measured transcriptomic, metabolomic and signaling changes driven by PI3Kis in cisplatin-resistant HNSCCs *in vitro* and tested efficacy *in vivo* in subcutaneous, orthotopic and metastatic xenograft models using immunodeficient and humanized murine models of HNSCC coupled with spatial transcriptomics.

**Results:** The PI3K pathway is activated in Nrf2-driven cisplatin-resistant HNSCC and is suitable for blockade as demonstrated in an *in vivo* shRNA screen. The PI3Ki gedatolisib inhibits cisplatin-resistant HNSCC proliferation, induces G2M arrest and potentiates cisplatin effectiveness through activation of autophagy, senescence and disruption of fatty acid metabolism. Gedatolisib suppresses HNSCC tumor growth in orthotopic and metastatic settings and demonstrates profound anti-tumor activity in humanized murine models of HNSCC, coupled with a reduction in hypoxia-rich regions and reduced infiltration by regulatory T lymphocytes.

**Conclusion:** Our findings emphasize the critical role of the PI3K-AKT-mTOR pathway in cisplatin-resistant HNSCC and highlight the therapeutic potential of PI3K inhibitors. Gedatolisib induced metabolic regulation and substantial re-sensitization of resistant cells to cisplatin, positioning it as a promising candidate for combination therapies aimed at overcoming primary chemo-radiation failure in HNSCC.

**Statement of translational relevance:** Cisplatin resistance, whether intrinsic or acquired, translates to treatment failure and nearly universal death in head and neck squamous cell carcinoma (HNSCC). However, the development of effective systemic regimens for cisplatin-resistant HNSCC has not yet been successful. Here, we present, for the first time, a mechanistic, biomarker-informed strategy for effective targeting of the PI3Kinase pathway in cisplatin-resistant HNSCC with substantial anti-tumor activity in both orthotopic and metastatic models, which may be capable of bypassing or reversing cisplatin resistance in this disease.

## Introduction

Head and neck squamous cell carcinoma (HNSCC) is the seventh leading cause of cancer-related mortality worldwide. In the recurrent/metastatic setting, the disease is nearly uniformly fatal due to a lack of effective systemic treatments(1,2). This oncologic reality is driven by the genomic landscape of the disease, defined by our group and others a decade ago (3–5). Most recurring genomic alterations identified in HNSCC are tumor suppressors and not directly targetable (5,6). Activating *HRAS* mutations are found in 6% of HNSCC tumors, but specific inhibitors akin to those targeting mutant (mut) *KRAS* are only in early stages of development, as is targeting Cyclin D1 amplifications with CDK4/6 inhibitors. In addition, although immune checkpoint inhibitors (ICIs) have been approved for recurrent metastatic HNSCC, their efficacy is generally low (<20%) and true cures are uncommon (∼7%) (7). Thus, cisplatin remains the mainstay systemic agent for the definitive treatment of HNSCC, with proven superiority over targeted agents (e.g., cetuximab), and is a critical component of systemic treatment (combined with ICIs) for recurrent/metastatic disease in patients with low combined positive scores (CPS) (8). Unfortunately, continued exposure to cisplatin can drive the development of acquired resistance, and some HNSCC tumors appear to be predisposed to limited responsiveness through *a priori* intrinsic resistance (9–11). This phenotype, now shown by us and others to be driven by hyperactivation of the Nrf2 pathway and supported by metabolic shifts favoring an enhanced reductive state, appears at first glance nearly untargetable and strongly coincides with high metastatic potential and suppressed anti-tumor immunity (10–12).

Activating *PIK3CA* mutations are present in approximately 18% of HNSCCs (4) and up to 30% of other solid tumors (e.g., breast cancer, endometrial cancer), and multiple clinically viable PI3K inhibitors (PI3Ki) exist. Indeed, PI3K/AKT/mTOR pathway inhibitors have been tested in a greater number of phase II/III trials than any other targeted therapy (13) owing to the frequent dysregulation of this pathway in cancer. Despite their clinical potential, PI3Kis have demonstrated only modest efficacy in the treatment of cancers harboring activating *PIK3CA* mutations, and the presence of *PIK3CA* mutations is not consistently predictive of drug response (14). Two recent discoveries, however, may improve the landscape of PI3K inhibition in squamous cell carcinomas such as HNSCC. First, our group discovered that *NOTCH1*-mutant HNSCCs are extremely sensitive to PI3Ki and undergo cell death rather than just growth arrest, in contrast to HNSCC tumors harboring *PIK3CA* mutations (15–18). Second, a novel potent and broad-acting PI3K/mTOR inhibitor, gedatolisib, has shown promising results in advanced breast cancer trials, with a very low incidence of hyperglycemia (6% grade 3/4) or pneumonitis (3%) (19). In this study, we sought to test the hypothesis that PI3K inhibition can overcome chemo-radiation resistance driven by Nrf2 hyperactivation, using our previous finding that *NOTCH1*-mutant head and neck squamous cell carcinoma responds favorably to this therapeutic approach. We leveraged HNSCC models we have built over the last 7 years, with demonstrated *in vitro* and *in vivo* cisplatin resistance that is functionally dependent on Nrf2 activation and associated with high rates of both cervical and distant metastasis.(10,11,20–23)

## Materials and Methods

### Cell lines

HN30 (RRID:CVCL_5525), HN31 (RRID:CVCL_5526), PCI-13(RRID:CVCL_C182), SCC152 (RRID:CVCL_C058), SCC154 (RRID:CVCL_2230), UDSCC2 (RRID:CVCL_E325), UMSCC47 (RRID:CVCL_7759) HN30R8, HN31P10, and PCI-13-4E were maintained in complete DMEM supplemented with 10% fetal bovine serum (FBS), 1% vitamins, sodium pyruvate, penicillin-streptomycin, L-glutamine, and non-essential amino acids (11,22). Cells were routinely passaged in complete growth medium containing 0.2% Myco-Zap (Lonza) to prevent mycoplasma contamination. Authentication of the cell lines was confirmed by Short Tandem Repeat (STR) profiling every 3 months. The cisplatin-resistant variants HN30R8, HN31P10 and PCI134E were maintained in cisplatin as previously described (10,11).

### Western blot

Western blot analysis was performed as previously described (12). Briefly, cells were lysed in RIPA lysis extraction buffer (Thermo Fisher Scientific Cat# 89900, Waltham, MA, USA) supplemented with a protease and phosphatase inhibitor cocktail (Thermo Fisher Scientific #78442, Waltham, MA, USA), and protein concentrations were determined using the BCA Protein Assay Kit (Thermo Fisher Scientific Cat# 23225, Waltham, MA, USA). Equal amounts of protein were separated by SDS-PAGE and transferred onto polyvinylidene fluoride (PVDF) membranes (Millipore Cat# IPVH00010, Burlington, MA, USA). Membranes were blocked with 5% skim milk in TBS-T, incubated overnight at 4°C with primary antibodies **(supplementary table 1)**, washed, and treated with HRP-conjugated secondary antibodies **(supplementary table 1)**. Protein bands were visualized using an enhanced chemiluminescence (ECL) reagent (BioRad Cat# 1705061, Hercules, CA, USA) and imaged with a ChemiDoc imaging system (Bio-Rad Laboratories, Hercules, CA, USA). Densitometry analysis was performed using Image Studio Lite v2 (LI-COR Biosciences, Lincoln, NE, USA). The results were normalized to β-actin as a loading control.

### Proliferation assay

A 96-well plate was seeded at 10,000 cells/well for four cell lines: HN30, HN30R8, HN31, and HN31P10. After a 24-hour incubation period, the cells were treated with varying concentrations of copanlisib (MedChemExpress Cat# HY-15346A, Monmouth Junction, NJ, USA) or gedatolisib (Selleckchem Cat# S2628, Houston, TX, USA). After 72 h of treatment, cell proliferation was assessed using either the Hoechst or resazurin assay. For the Hoechst assay, media were aspirated from each well, followed by two cycles of freeze-thawing; cells were initially frozen at -80°C, thawed, and then refrozen at -80°C after adding 100 µL of water to each well. After the second thaw, 100 µL of TNE buffer containing Hoechst dye (Thermo Fisher Scientific, Cat# 62249, Waltham, MA, USA) (20 µL per 10 mL of TNE buffer) was added to each well. Alternatively, for the resazurin assay, 1:10 resazurin reagent (Cat# AR002; R&D Systems, Minneapolis, MN, USA) was added to the cell culture medium. Both assays were performed using fluorescent readings. Data were normalized to day 0 and GraphPad Prism software (GraphPad Software, Inc., San Diego, CA, USA) was used to plot the percentage of cell viability and calculate the IC_50_ for each cell line.

### Xenograft mouse models of oral cancer and therapy

All animal experiments were conducted in compliance with protocols approved by the Institutional Animal Care and Use Committee (IACUC) at Baylor College of Medicine. In this study we utilized six-week-old female BALB/C nude mice and 14-week -old NOD humanized mice (HLA-A matched to human cell line) (Taconic Biosciences, Hudson, NY, USA) for subcutaneous and orthotopic xenograft generation. For the subcutaneous model, each mouse received 1×10^6 HN30R8 cells in 100 µL PBS into the flank. For the orthotopic model, each mouse received 5×10^4 HN30R8 cells in 50 µL of PBS into the oral tongue. Once tumors formed, mice were randomly assigned to two groups: one receiving either copanlisib or gedatolisib treatment and the control group receiving saline. Each treatment group consisted of 5–10 mice. Copanlisib was administered intraperitoneally (I.P.) for 2 consecutive days, followed by a 1-day recovery period. Initially, the treatment dose for athymic nude mice was 14 mg/kg, which was later reduced to 10 mg/kg secondary to observed weight loss. For gedatolisib treatment, clinical-grade gedatolisib (Celcuity Inc., Minneapolis, MN, USA) was administered at a dose of 20 mg/kg in athymic nude mice and 16 mg/kg in humanized mice via tail vein injection every three days. Tumor volume was measured every three days by an investigator blinded to the treatment groups, using the formula *V*=(*L*×*W*^2^)/2, where *L* is the tumor length and *W* is the tumor width, both in millimeters. Following humane euthanasia, tumors were removed for analysis. The condition of all animals was closely monitored throughout the study, with particular attention to weight loss and any other adverse drug effects. For the metastatic model, six-week-old female *BALB/c* nude mice were injected with 50,000 HN30R8-luciferase cells via tail vein injection. Bioluminescence images were captured on day 0 and weekly thereafter to monitor tumor signal development. Once the bioluminescent signals reached a consistent region of interest (ROI) of approximately 100,000 photon counts, the mice were divided into two groups: a control group and a gedatolisib-treated group. The gedatolisib-treated group received 20 mg/kg of gedatolisib via tail vein injection twice a week, while the control group received saline.

### *In vivo* shRNA pooled screens and data analysis

*In vivo* shRNA screens were conducted as we previously published (PMID 34732714). Briefly, *NOTCH1* mutant HNSCC cell lines HN31, UMSCC22A, and UMSCC47 (HPV-associated) were infected *in vitro* with a pooled shRNA lentiviral library targeting 195 druggable genes (average 10 hairpins per gene) After a two-day puromycin selection, cells were minimally expanded and either frozen down for time zero reference points or injected subcutaneously (4 million per animal) into mice. After tumor formation, mice were randomized to receive regimens of either fractionated radiation or carboplatin at doses predetermined to inhibit tumor growth by 25% (ED25) or no treatment at all. Genomic DNA harvested from tumors reaching approximately 500mm^3^ was used to amplify vector sequences which were then prepared for next generation sequencing. Count data was analyzed using the siRNA-activity (RSA) algorithm to derive P-values and log2 fold changes for each targeted gene according to our published methods (PMID 34732714).

### Humanized mice flow cytometry staining

At the time of sacrifice, blood samples were collected via an intracardiac puncture post-mortem. Circulating lymphocytes were enriched by layering blood over Lymphocyte Separation Buffer (Corning) and centrifuging at 850 × *g* for 20 min. Spleens were harvested, mechanically disrupted through a 70 µm cell strainer, and treated with RBC Lysis Buffer (Cat# SKU TNB-4300; Tonbo Biosciences, San Diego, CA, USA) according to the manufacturer’s protocol to isolate white blood cells. The resulting suspensions were filtered to obtain single-cell suspensions for flow cytometry. Cell suspensions were stained with Ghost Dye™ UV 450 (Tonbo Biosciences, Cat# SKU 13-0868, San Diego, CA, USA) to identify dead cells and subsequently incubated with Purified Anti-Mouse CD16/CD32 Fc Shield (Tonbo Biosciences Cat# SKU 70-0161, San Diego, CA, USA). Cells were then washed and labeled with fluorescently conjugated surface antibodies **(supplementary table 2)** in brilliant stain buffer (Invitrogen, Cat# 00-4409-42). Staining was performed at 4°C for 30 min. Prior to intracellular marker staining, cells were fixed and permeabilized using the Foxp3/Transcription Factor Staining Buffer Kit (Tonbo Biosciences, Cat# SKU TNB-0607, San Diego, CA, USA) following the manufacturer’s instructions. Data acquisition was conducted on a 5-laser Cytek Aurora flow cytometer, spectrally unmixed with SpectroFlo (Cytek Biosciences) and the resulting data were analyzed using FlowJo v10.10 and GraphPad Prism 10 for visualization and comprehensive statistical evaluation.

### Tissue analysis

Tissue slides were processed for immunohistochemistry (IHC) by first deparaffinizing in xylene, followed by rehydration through a graded series of ethanol solutions diluted in water. To inhibit endogenous peroxidase activity, slides were treated with 3% hydrogen peroxide for 10 min. Antigen retrieval was performed using heat-induced epitope retrieval for 20 min to facilitate unmasking of target proteins. Manual IHC staining was conducted for markers including CD3 (clone LN10) (Leica Biosystems Cat# PA0122, RRID:AB_3073619, Buffalo Grove, IL, USA), CD4 (clone 4B12) Leica Biosystems Cat# PA0371, RRID:AB_10554438,, Buffalo Grove, IL, USA), CD8 (clone 4B11) (Leica Biosystems Cat# PA0183, RRID:AB_10555292, Buffalo Grove, IL, USA) p40 (clone clone BC28) (Leica Biosystems Cat# PA0163, RRID:AB_3073535, Buffalo Grove, IL, USA), 4B12, and 4B11 respectively, sourced from Leica Biosystems, Buffalo Grove, IL, USA), and pan-cytokeratin (panCK, Clone E6S1S) (Cell Signaling Cat#83957, Danvers, MA, USA). Detection was achieved using the Bond Polymer Refine Detection system (Leica Biosystems, Buffalo Grove, IL, USA), utilizing horseradish peroxidase (HRP) to catalyze the chromogenic reaction. The 3,3′-diaminobenzidine (DAB) Chromogen Kit (Leica Biosystems, Buffalo Grove, IL, USA) was used to visualize the target proteins, producing a brown stain that indicated positive expression. Appropriate positive and negative controls were included to ensure specificity and reliability of the staining results. Stained slides were imaged using a Nikon Eclipse slide-scanning microscope equipped with Plan Apo objectives. Images were captured at magnifications of 10X or 20X, with numerical apertures of 0.45 and 0.20, respectively. This approach allows for high-resolution visualization of cellular and tissue morphology, facilitating a detailed analysis of protein expression across samples.

### Steady state unbiased metabolomics

To assess the metabolic impact of copanlisib, cells were seeded at a density of 3 million cells per 15 cm dish and pre-treated with 10 µM cisplatin for 48 h. On the second day, the cells were gently washed twice with 25 mL of PBS and once with 15 mL of fresh medium to remove the residual treatment. Subsequently, 25 mL fresh growth medium was added. The treatment group received 30 nM (IC_50_) copanlisib (MedChemExpress Cat# HY-15346, Monmouth Junction, NJ, USA), while the control group was administered an equal volume of saline. The cells were incubated for 24 h under these conditions before harvesting for metabolomic analysis. Similarly, to study the effects of gedatolisib, 3 million cells were plated per 15 cm dish and allowed to recover in the presence of 10 µM cisplatin for 48 h. Cells were then washed with PBS and fresh medium, after which 25 mL of growth medium was added. The treated group received 20 nM (IC_50_) gedatolisib (Selleckchem Cat#S2628, Houston, TX, USA), and the control group was exposed to the same volume of DMSO. Following a 24-hour incubation, the cells were collected for metabolomic profiling. For untargeted metabolomics, cells were subjected to a freeze-thaw cycle and homogenized using needle sonication in a methanol/water mixture (1:1). Three volumes of methanol/acetonitrile (1:1) were added, and the samples were vortexed for 5 min and then kept at -20°C for 10 min to precipitate proteins. After centrifugation at 15,000 rpm for 10 min at 4°C, the supernatants were evaporated using GeneVac EZ-2 Plus SpeedVac (SP Scientific). The dried residues were reconstituted in 100 µL methanol/water (50:50 v/v). Quality control (QC) samples were prepared by pooling 20 µL of each supernatant with aliquots used for QC during analysis. Chromatographic separation utilized A Thermo Scientific Vanquish Horizon UHPLC system, with two columns: a Waters ACQUITY HSS T3 column (1.8 µm, 2.1 mm x 150 mm) for reverse-phase (RP) separation, and a Waters ACQUITY BEH amide column (1.7 µm, 2.1 mm x 150 mm) for hydrophilic interaction liquid chromatography (HILIC). The RP method employed a gradient from 99% mobile phase A (0.1% formic acid in water) to 95% mobile phase B (0.1% formic acid in methanol) for 21 min. HILIC separation was performed using gradients of acetonitrile and water with 0.1% formic acid and 10 mM ammonium formate. Both column methods were run at 50°C with a flow rate of 300 µL/min and a 2 µL injection volume. Data acquisition was performed using a Thermo Scientific Orbitrap IQ-X Tribrid Mass Spectrometer, with settings adjusted for the positive (3500 V spray voltage) and negative (2500 V spray voltage) modes. The vaporizer and ion transfer tube temperatures were set at 300°C, whereas the auxiliary gas heater was set at 350°C. The sheath, auxiliary, and sweep gases were set at 40, 8, and 1 arbitrary units, respectively. Full MS scans were carried out with a resolution of 120,000, spanning a mass range of 70–900 m/z, using quadrupole isolation and automatic gain control (AGC) parameters of 25% and 1.0e5, respectively. Further compound analysis was conducted via data-dependent MS/MS using high-energy collisional dissociation (HCD) at collision energies of 30%, 50%, and 150%. Data collection utilized a resolution of 30,000, with normalized AGC targeting 100% and an absolute AGC value of 5.0e5 capped at a 50 ms injection time. Data processing was performed using Compound Discoverer (v3.3.3.2, Thermo Fisher Scientific), enabling peak detection, integration, and identification via the mzCloud and NIST 2020 high-resolution mass spectral library, in-house library of 600 compounds, and Human Metabolome Database (HMDB). The QC samples were run every 10 injections for consistency. Statistical analyses involved log2 transformation and normalization using the median interquartile range (IQR). Differential metabolites were identified using Student’s t-test with a false discovery rate (FDR < 0.25) controlled by the Benjamini-Hochberg method. Metabolites with p-values <0.05 were further analyzed, excluding non-biological compounds based on HMDB were further analyzed. The identified biological metabolites were visualized through double hierarchical clustering using the Morpheus platform, with z-score normalization applied to the data.

### RNA-Sequencing

Total RNA from biological replicates was extracted using a RNeasy Mini Kit (Qiagen Cat#74104, Germantown, MD, USA), according to the manufacturer’s instructions. RNA was subjected to whole transcriptomic sequencing (Psomagen Inc.) to produce RSEM counts and effective gene sizes. For statistical analysis, FPKM-UQ values were calculated and log-transformed as Log2(X + 0.01). Single-sample gene set enrichment analysis (ssGSEA) was conducted in linear space using the Broad Institute’s Gene Pattern platform. Additionally, RNA-seq RSEM data from TCGA, harmonized through the Toil open-source data portal, were normalized as previously described. For differential gene expression, only genes with an average log2 FPKM value of ≥ 2 in at least one group (treatment or control) were included. Differential expression analysis was performed using JMP 13 software, with t-tests for each gene and Benjamini-Hochberg correction (FDR < 0.1) for adjusted P-values. Fold changes were calculated as the ratio of geometric means (2^[log-fold change]). Previously published and validated Nrf2 pathway gene lists were used for ssGSEA scoring. Gene expression data cross-correlation coefficients were analyzed via two-way hierarchical clustering to identify modules with similar expression patterns, validated against independent datasets, or through orthogonal methods. Our in-house MATLAB script, based on resampling techniques for consensus hierarchical clustering, was employed using Ward’s linkage (12,24). This script is publicly available at GitHub: https://github.com/aif33/Hierarchical-two-way-agglomerative-consensus-clustering.

### Xenium Spatial Transcriptomics

Xenium slides were prepared following the Xenium In Situ for Fresh Frozen Tissues User Guide CG000579 from 10x Genomics. Briefly, fresh tumor samples were placed into Optimal Cutting Temperature (OCT) compound and flash-frozen in an isopentane/dry ice bath. 10 µm-thick tissue sections were placed onto the Xenium slide capture area and stored at -80 °C for up to one week. Slides were subsequently fixed with paraformaldehyde and permeabilized using MeOH as per the manufacturer’s protocol (10x Genomics User Guide CG000581). Following fixation and permeabilization, the tissue sections were processed and stained according to the manufacturer’s protocol (10x Genomics User Guide CG000760). Briefly, ssDNA probes were bound to RNA, followed by ligation and rolling circle amplification. Finally, cells were stained for cell segmentation with 10x Genomics’ Xenium In Situ Cell Segmentation Kit. The probe set consisted of the predesigned “Xenium Prime 5K Human Pan Tissue & Pathways Panel” and a custom add-on panel of 45 probes targeting both human and mouse transcripts **(supplementary table 3)**. The prepared slides were run on the Xenium Analyzer at the Baylor College of Medicine Single Cell Genomics Core, using v3.0 analysis, with capture regions selected to cover each tumor section. Immediately following the run, hematoxylin and eosin (H&E) staining was performed on the tissue sections using the manufacturer’s protocol (10x Genomics User Guide CG000613). H&E images were acquired with a Keyence BZ-X810 microscope. Xenium data was processed using the Seurat package in R and visualized with Seurat, ggplot2, and Xenium Explorer (10x Genomics).(25,26) For gene expression analysis, count data from each tissue were combined, log-normalized, and scaled in Seurat. Each cell was annotated as of mouse or human origin based on the species of the highest-expressed gene. For human cells, Uniform Manifold Approximation and Projection (UMAP)(27) and nearest-neighbor graph construction were performed with 16 principal components and cluster determination with a resolution of 0.3. Differential gene expression was calculated in Seurat with the FindMarkers function. Gene set enrichment analysis was performed with the fgsea package using sign(FC) * -log_10_(p_adj_) as the ranking statistic, where FC indicates the log_2_ fold change and p_adj_ the Bonferroni-adjusted p-value of the relevant differential gene expression calculation. (28)

### Statistical Analysis

For cell cycle experiments with multiple treatments, the logit transformation was applied to percentages and differences compared to control were determined with an ANOVA and post-hoc Dunett’s test with further family wise corrections for multiple testing within each treatment using an inhouse Matlab script we previously described. Where only two groups were compared, multiple T-testing was used with a Benjamini-Hochberg correction (FDR =0.1). For the cell growth experiments combining cisplatin and gedatolisib, background subtracted fluorescent values were considered as the response variable and a three-way ANOVA model was evaluated with main effects treatment, dose, and regimen. Post-hoc contrast between groups were analyzed by Tukey’s Honestly Significant Difference test (FDR =0.05).

## Results

### Comparative transcriptomic analysis reveals Nrf2 and PI3K-AKT-mTOR pathway alterations in cisplatin-resistant HNSCC

We conducted western blot analyses to assess the expression levels of keap1, nrf2, gpx2, and p53 in two isogenic pairs of HNSCC cell lines and their cisplatin-resistant derivatives. Cisplatin-resistant cells exhibited a marked reduction in keap1, accompanied by increased nrf2 levels compared to their parental counterparts **(figure S1A)**. HN31 cells displayed elevated p53 protein levels, consistent with the mutational status of *TP53* compared to protein levels of the functional p53 protein in HN30 **(figure S1A)**. RNAseq was used to examine the landscape of differentially expressed genes (DEGs) between CDDP resistant cell lines and their matched parental cell lines. Initially we determined the overlap between DEGs and those from a 138-gene NRF2 signature we previously derived (12) that includes known upregulated NRF2 targets. A total of 57 NRF2-dependent genes were commonly upregulated greater than 1.3-fold in CDDP resistant cell lines derived from both backgrounds (**figure S1B; supplementary tables 4 -5**), while only 7 of the NRF2-regualted genes were commonly downregulated (**figure S1D**; **supplementary tables 4-5**), indicating a preponderance of NRF2 activation. Analysis of the additional 1202 genes commonly upregulated in the cisplatin resistant cells (**figure S1C**; **supplementary tables 6-7**) identified highly significant enrichment for multiple Gene Ontology hallmark pathways (**figure S1C**; **supplementary table 8)**, including PI3K-AKT-mTOR. Although many genes were also downregulated in cisplatin resistant HN31P10, that was not true for HN30R8 and only 8 total genes were commonly downregulated for both (**figure S1E; supplementary tables 6-7**) with no pathway enrichment found. Analysis of a previously completed *in vivo* shRNA screen **(figure S2A-C)** identified the PI3K-AKT-mTOR pathway as a suitable target for therapeutic intervention across various genomic backgrounds regardless of association with the human papillomavirus (HPV), with increased dependency in the presence of carboplatin for at least two cell lines examined **(figure S2D).**

### Evaluating the efficacy of PI3K inhibition in cisplatin-resistant HNSCC

To test whether inferred reliance on the PI3K-AKT-mTOR pathway translated into targeting sensitivity, we initially treated the resistant cell lines with the pan-PI3K inhibitor copanlisib, which demonstrated comparable IC_50_ values for both cisplatin-resistant and parental cell lines (32nM-HN30R8, 43nM-HN31P10, 50nM-HN30, 34nM-HN31 cell lines) **(figure S3A-B)**. Subcutaneous and orthotopic xenograft models of the HN30R8 cell line showed a significant delay in tumor growth when treated with 10 mg/kg copanlisib **(figures S3C-G)**. However, the treatment was associated with dose-limiting toxicity, as evidenced by weight loss in the treated mice **(figures S3D, S3G)**. Interestingly, mice with orthotopic tumors showed greater weight loss, possibly due to the impact of the primary tumor on oral intake. We next evaluated a PI3Ki with an expectedly lower toxicity profile and dual inhibition of both PI3K and mTOR—gedatolisib. Gedatolisib IC_50_ values were again comparable between cisplatin-resistant lines and their parental counterparts (22nM -HN30R8, 18nM-HN31P10, 17nM -HN30, and 9nM-HN31 cell lines) **(figure 1A-B)**. *In vivo* experiments using both an orthotopic model and a metastatic model demonstrated significant tumor growth suppression in the gedatolisib-treated group (20 mg/kg), as evidenced by tumor growth delay, decreased IVIS signal, and gross anatomical assessment **(figures 1C-G, S5)**. Unlike copanlisib, gedatolisib-treated mice did not exhibit measurable weight loss or other dose limiting toxicity **(figure 1D)**, highlighting its potential as a more tolerable therapeutic option. To expand the potential applicability of gedatolisib, we demonstrated *in vitro* efficacy at nM concentrations against both HPV-associated and HPV-independent HNSCC cell lines, and a 3^rd^ parental-cisplatin-resistant isogenic pair generated in the PCI-13 background. Of note, gedatolisib demonstrated good activity across cell lines that spanned a spectrum of sensitivity and resistance when previously tested against other conventional Pi3Kis **(figure S4)**.

**Figure 1.**
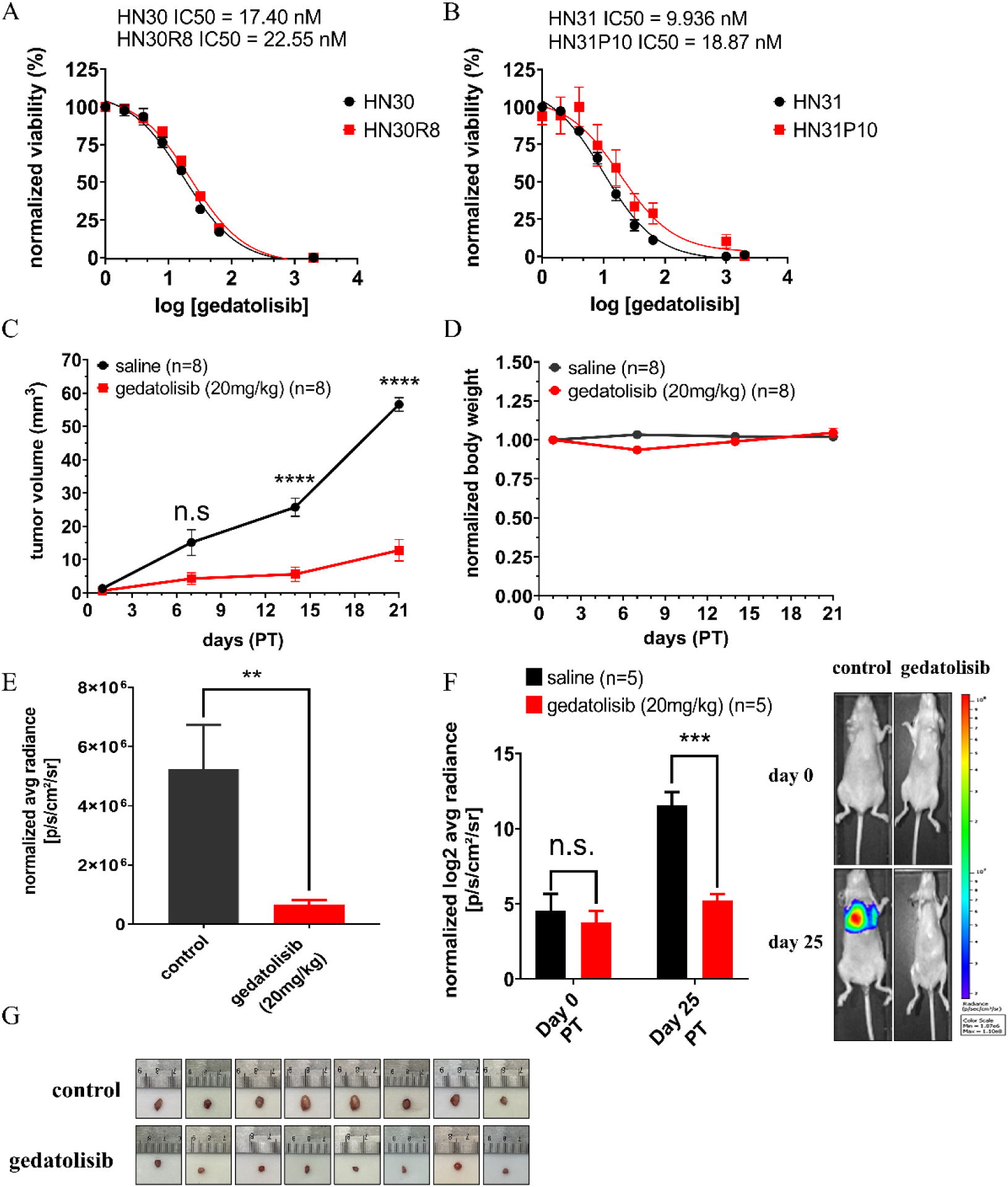
Gedatolisib suppresses HNSCC cisplatin-resistant tumor growth. **(A-B)** HNSCC cell lines were treated with gedatolisib for 72 hours and cell number and viability were assessed using a Resazurin assay. The sigmoid graph plots normalized cell viability (y-axis) against the logarithm of gedatolisib concentration (x-axis) to determine the IC_50_. Black represents parental cell lines, while red represents cisplatin-resistant cell lines. **(C)** *In vivo* data from orthotopic xenograft models using HN30R8 cisplatin-resistant cell lines treated with 20 mg/kg of gedatolisib via tail vein (IV) administration every 3 days. The y-axis represents tumor volume (mm²), and the x-axis represents days post-injection following tumor visualization. Black data points indicate the control group, while red data points represent the gedatolisib-treated group. Data are presented as the mean ± standard error of the mean (SEM). P-values were calculated using a two-tailed Student’s t-test (* P ≤ 0.05, ** P ≤ 0.01, *** P ≤ 0.001, **** P ≤ 0.0001). **(D)** Normalized body weight representation of *in vivo* orthotopic xenograft models treated with 20mg/kg of gedatolisib. The y-axis represents body weight, and the x-axis denotes days post-injection following tumor visualization. Black data points indicate the control group, while red data points represent the gedatolisib-treated group. **(E)** IVIS signal showing luciferase activity in mice bearing HN30R8 tumors, at day 21 post-treatment. The bar graph displays luminescence signals for both control and treated groups. The y-axis represents normalized radiance to day 0, and the x-axis indicates the mouse groups. Data are presented as the mean ± standard error of the mean (SEM). P-values were determined using a two-tailed Student’s t-test (ns, not significant; * P ≤ 0.05). **(F)** Luciferase activity in mice bearing metastatic HN30R8 tumors, at day 25 post-treatment initiation (PT). The y-axis represents average radiance normalized to day 0, and the x-axis indicates day 0 and day 25 post treatment initiation (PT). Data are presented as the mean ± standard error of the mean (SEM). P-values were determined using a two-tailed Student’s t-test (ns, not significant; * P ≤ 0.05, ** P ≤ 0.01, *** P ≤ 0.001). **(G)** Images of extracted tumors from orthotopic model.

### Mechanistic analysis of acute and delayed effects of PI3Ki in cisplatin-resistant HNSCC

To assess gedatolisib’s putative mechanism of action, HN30R8 and HN31P10 cells were treated with PI3Kis for 6h for acute and 48h for delayed effect assessment. Acute exposure to copanlisib did not result in significant inhibition of phosphorylated Akt, although it effectively reduced the phosphorylation of downstream targets such as p70S6, p4EBP1, and pS6 **(figure S6A)**. In contrast, gedatolisib rapidly inhibited Akt phosphorylation in both cell lines, along with reductions in p70S6, p4EBP1, pS6, and pmTOR, and a slight decrease in total mTOR levels **(figure 2A)**. At 48 hours post exposure, copanlisib moderately inhibited phosphorylation of Akt, Gsk3β, 4EBP1, and S6 **(figure S6B),** while gedatolisib **(figure 2B)** exhibited a more comprehensive and sustained effect on PI3K signaling. To further elucidate the mechanism of action of gedatolisib in cisplatin-resistant cell lines, we subjected HN30R8 and HN31P10 to transcriptomic analysis following drug treatment for 24 and 48 h **(figures 3A, S7A-D; supplementary tables 9-16)**. At 24 h, gedatolisib predominantly downregulated genes involved in cell cycle regulation and cell division **(figure S7A; supplementary tables 9,11)**. Interestingly, after 48 h, the impact of gedatolisib shifted towards the downregulation of metabolic processes **(figure S7B; supplementary tables 10,12)**. In terms of upregulated genes, at 24 h, gedatolisib enhanced transcripts related to the cellular stress response and morphogenesis **(figure S7C; supplementary table 14)**. After 48 h, sustained upregulation of genes associated with stress response was observed **(figure S7D; supplementary table 15)**. We further conducted a comprehensive analysis of the common genes across both time points by examining both the upregulated and downregulated transcripts. This analysis highlighted significant downregulation of metabolic pathways, including the key Nrf2 targets *AKR1C1*, *AKR1C2, AKR1B10*, and *AKR1B15,* known to mediate cisplatin resistance **(figure 3A; supplementary table 13)**. Autophagy genes that were consistently upregulated at both timepoints included *ATG16L2*, *CTSF*, and *CREB3L4* **(figure S7E; supplementary table 16)**.

**Figure 2.**
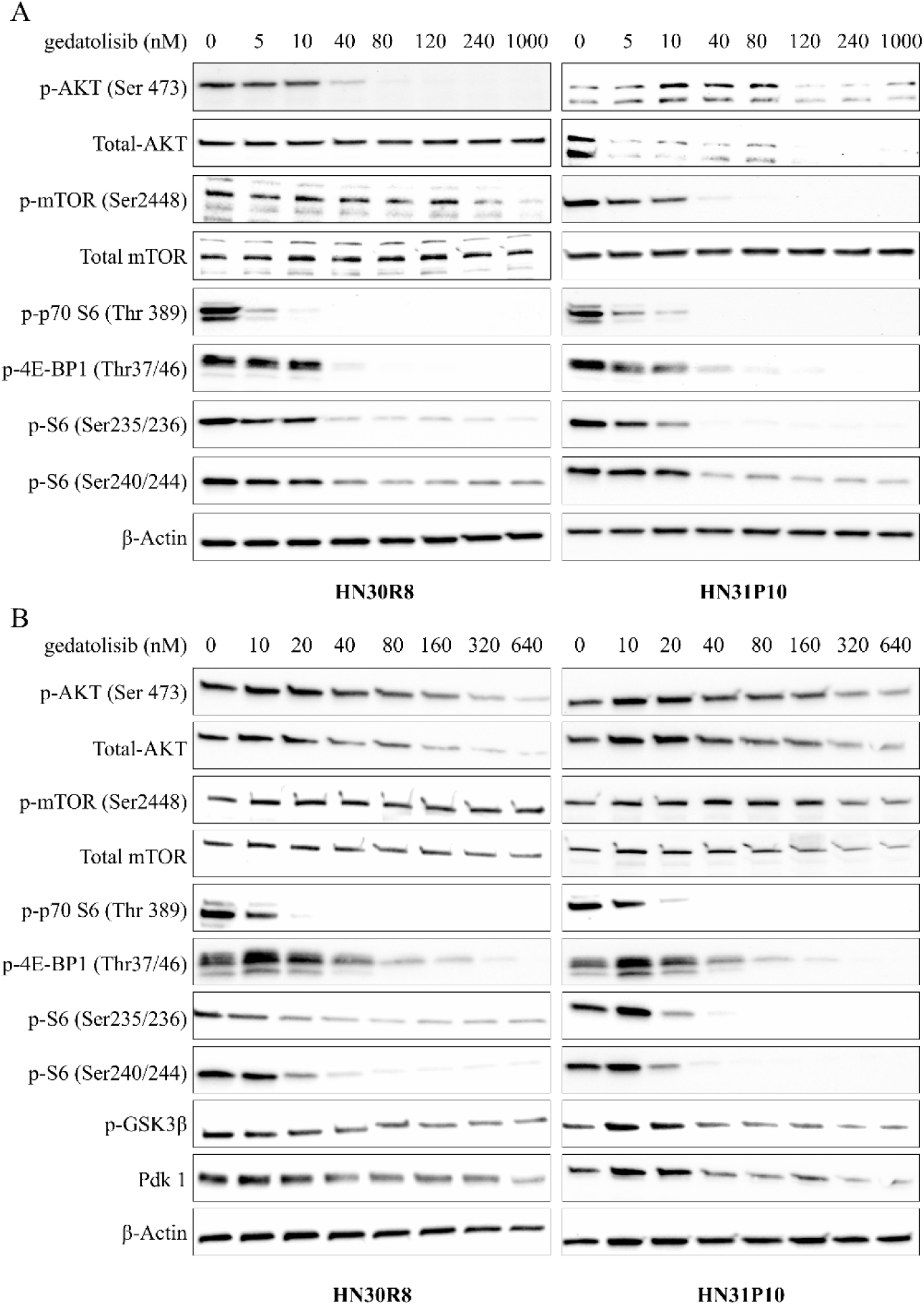
Gedatolisib inhibits PI3K pathway activity. **(A)** Representative Western blot data showing selected PI3K downstream target proteins. Cisplatin-resistant cell lines, HN30R8 (left panel) and HN31P10 (right panel), were treated with gedatolisib and/or control for 6 hours at the indicated drug concentrations (nM). β-Actin was used as a loading control. **(B)** Representative Western blot data showing selected PI3K downstream target proteins. Cisplatin-resistant cell lines, HN30R8 (left panel) and HN31P10 (right panel), were treated with gedatolisib and/or control for 48 hours at the indicated drug concentrations (nM). β-actin was used as a loading control.

**Figure 3.**
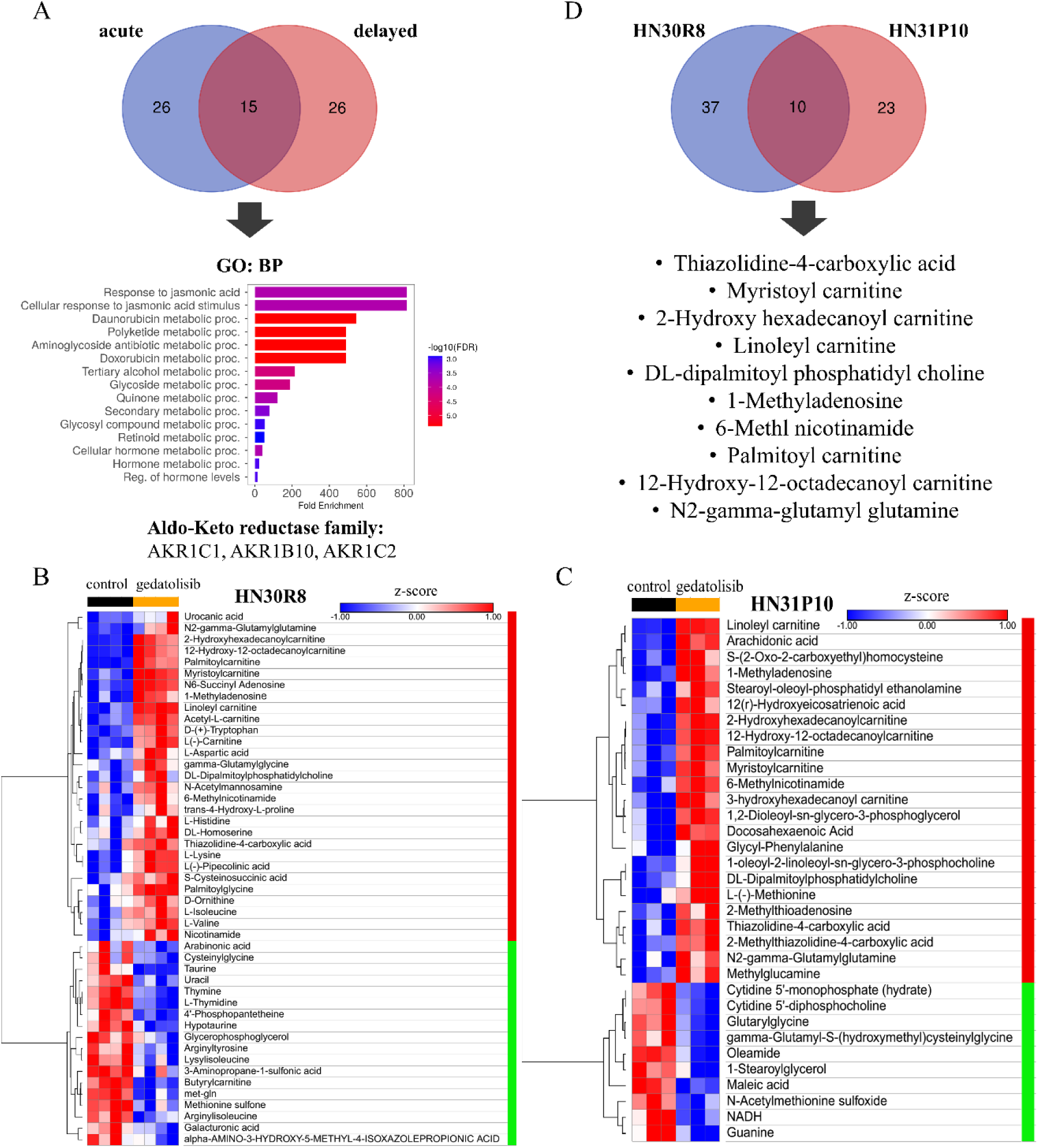
Gedatolisib effects on steady-state metabolism in cisplatin-resistant HNSCC. **(A)** Differential gene expression (DGE) analysis showing downregulated overlapping genes of acute (24 hours) and delayed (48 hours) time points of gedatolisib treatment, using a Venn diagram in HN30R8 and HN31P10 cisplatin-resistant cell lines. The bar graphs represent Gene Ontology (GO) analysis, with the y-axis showing enrichment pathways and the x-axis indicating fold enrichment. Data are sorted based on false discovery rate (FDR) with a cutoff ≤0.05 and fold enrichment. **(B-D)** Differential heatmap of metabolites in HN30R8 and HN31P10 cells treated with vehicle (control) or gedatolisib for 24 hours, along with a Venn diagram highlighting the common metabolites in both cell lines.

Given that gedatolisib may generate anti-tumor effects through metabolic regulation, we metabolically profiled its effects alongside copanlisib by treating HN30R8 and HN31P10 with their respective IC_50_ concentrations for 24 h. After treatment, steady-state, non-exposome related metabolites were analyzed. Copanlisib treatment consistently impacted four common metabolites: asparagine, threonine, gamma-glutamylglycine, and L-histidine **(figure S8A-C; supplementary tables 17-18)**. Gedatolisib treatment led to a greater metabolic effect, as shown by the larger number of differential metabolites *in common* across the two cell lines **(figure 3B; supplementary tables 19-20)** inclusive of: thiazolidine-4-carboxylic acid (T4CA), myristoyl carnitine, 2-hydroxy hexadecanoyl carnitine, linoleyl carnitine, DL-dipalmitoyl phosphatidyl choline, 1-methyladenosine, 6-methyl nicotinamide, palmitoyl carnitine, 12-hydroxy-12-octadecanoyl carnitine, and N2-gamma-glutamyl glutamine. These changes stemmed from increased numbers of altered metabolites in the individual CDDP-resistance cell lines (**figure 3C and 3D**). These findings suggest that gedatolisib may generate a broader metabolic disruption than copanlisib, although whether this is a primary or secondary mechanism of anti-tumor activity remains to be determined through deeper analyses of metabolic flux under exposure conditions.

### Exploring gedatolisib’s role in bypassing cisplatin resistance in HNSCC

Whereas copanlisib treatment of HNSCC reduced cell growth and induced limited cell death **(figures S9A-C, S10A-B)**, gedatolisib did not induce substantial cell death across cell lines **(figures S10C-D, S11A-B)** at 48 h as measured by increased cell permeability. At higher concentrations of gedatolisib, HN30R8 displayed a modest increase in cell death at 300 nM, while HN31P10 demonstrated approximately 15% cell death at 200 nM (**figure S11C-D).** At 24 h, gedatolisib caused a dose-dependent reduction in S-phase and an increase in G1-phase in HN30R8 (**figures 4A, S10G-H**) and HN31P10 (**figure S11E**), indicative of potential G1 cell cycle arrest. After 48 hours these differences subsided due to an increase in the sub-G1 phase population observed in both cell lines **(figures S11G-H, S10I-J)**. The observed cell death in both HNSCC cell lines was not primarily driven by caspase activation or PARP cleavage, which are the common indicators of apoptosis. Instead, there was a consistent and notable increase in autophagic activity across cell lines **(figures 4B, S11F)**.

**Figure 4.**
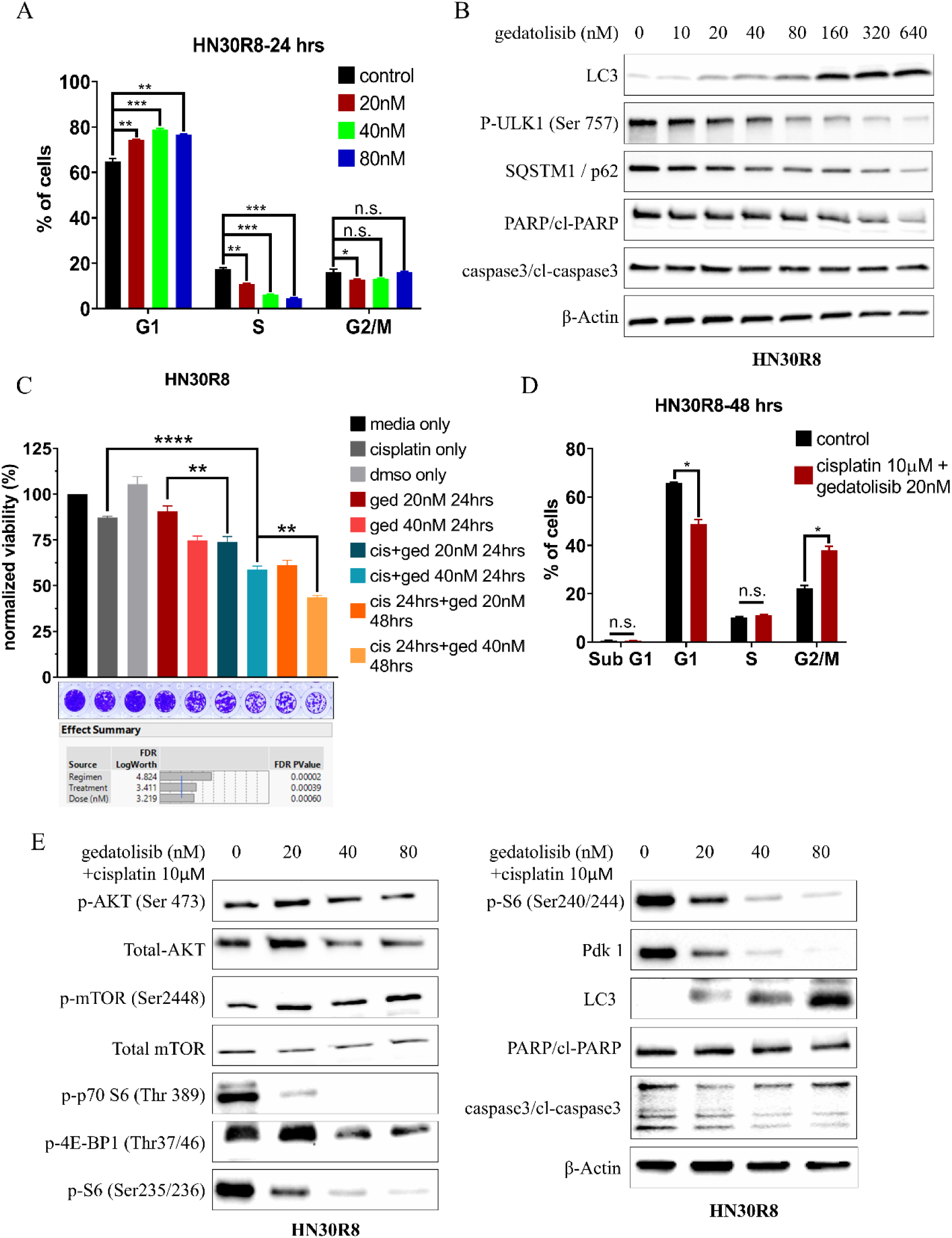
Gedatolisib induces cell cycle arrest and enhances sensitivity to cisplatin. **(A)** The bar graph panel represents flow cytometry cell cycle analysis for G1, S, and G2/M phases in HN30R8 exposed to gedatolisib for 24 hours. The y-axis represents the percentage of cells in each respective cell cycle phase, while the x-axis indicates the cell cycle phases, with each color corresponding to a specific concentration of gedatolisib. Data are presented as the mean ± standard error of the mean (SEM). P-values were calculated comparing treated groups to the control (0nM) using a one-way ANOVA Dunnet’s test (not significant (n.s.), * P ≤ 0.05, ** P ≤ 0.01, *** P ≤ 0.001). **(B)** Western blot panel for cell death determination in HN30R8 cell treated with gedatolisib for 48 hours. β-Actin was used as a loading control. **(C)** Clonogenic assay showing effects of gedatolisib and cisplatin combinations in HN30R8. The bar graph represents the cell viability of the cells obtained from same samples and normalized to day 0. The y-axis shows percentage of cell viability and x-axis represents the respective samples. Data are presented as the mean ± standard error of the mean (SEM). P-values were calculated using a three-way ANNOVA with Tukey’s test (** P ≤ 0.01, **** P ≤ 0.0001). **(D)** Flow cytometry cell cycle analysis bar graph showing sub G1, G1, S, and G2/M phases in HN30R8 treated with combination of gedatolisib and cisplatin for 48 hours. The y-axis represents the percentage of cells in each respective cell cycle phase, while the x-axis indicates the cell cycle phases. Black represents cisplatin alone, while red represents the combination of gedatolisib and cisplatin. Data are presented as the mean ± standard error of the mean (SEM). P-values were calculated using a one-way ANOVA with Dunnett’s test (not significant (n.s.), * P ≤ 0.05). **(E)** Western blot panel for HN30R8 treated with the combination of gedatolisib and cisplatin. β-actin was used as a loading control.

Given that gedatolisib significantly influenced cell metabolism and the role of the aldo-keto reductase family in cisplatin resistance (11,29), we investigated whether its addition could re-sensitize HN30R8 and HN31P10 to cisplatin. Cells were either pulsed with cisplatin, which was washed out before adding gedatolisib, or treated with both drugs simultaneously, and the inhibition of colony formation was compared to that of the single agents. As expected, cisplatin alone had minimal impact on HN30R8 or HN31P10. Gedatolisib alone provided some degree of suppression, but the most pronounced inhibition occurred when gedatolisib was combined with cisplatin in both HN30R8 and HN31P10 **(figures 4C-D, S12A-B; supplementary tables 21-22)**, regardless of whether cells were pretreated with cisplatin, or the two drugs were added simultaneously. When combined, gedatolisib and cisplatin led to a marked reduction in cell proliferation and G2/M cell cycle arrest **(figures 4D, S12B)**. Investigation of the signaling pathways revealed that the combination treatment had a nuanced effect on PI3K downstream kinases, with some variations in their modulation. However, a consistent and notable finding was the significant reduction in pdk1 levels, which plays a key role in cell survival and proliferation pathways. Additionally, there was a consistent increase in the expression of lc3b, and reduction of phosphorylated ulk1, and sqstm1/p62, markers of an active autophagic flux **(figures 4E, S12C-D)**. The presence of functional p53 allows the HN30 line to undergo senescence under conditions of stress. Here we found that gedatolisib, when combined with cisplatin, generated significant levels of cellular senescence (**figure S13 A-C).**

### Therapeutic potential of gedatolisib in humanized murine models

To further investigate the impact of gedatolisib on tumor progression, the tumor microenvironment, and immune cell infiltration, we employed a humanized orthotopic mouse model using cisplatin-resistant HN30R8. This humanized model reproduced the aggressive local growth pattern we previously described for the HN30R8 cisplatin-resistant model (10,11), with high rates of cervical metastasis and the development of spontaneous distant metastatic disease (lung) (**figure 5A**). Given that PI3K inhibitors have been reported to impair T cell activation and proliferation (30), we utilized a lower dose of gedatolisib (16 mg/kg) which nevertheless generated significant anti-tumor activity without loss of body weight **(figure 5B-C)**. We utilized flow cytometry analysis of blood and splenocytes (**figure S14**) to measure the effects of gedatolisib on systemic immunity. When analyzed as a fraction of human CD45^+^ cells, we detected modest but statistically significant reductions in plasma cells, central memory (CCR7^+^CD45RA^-^) T cells, and regulatory T cells (Tregs), and a statistically significant increase in early (CD56^+^CD16^−^) NK cells (**supplementary table 23**) but most tested immunocyte subsets were not affected by treatment to a significant degree **(figure S14)**.

**Figure 5.**
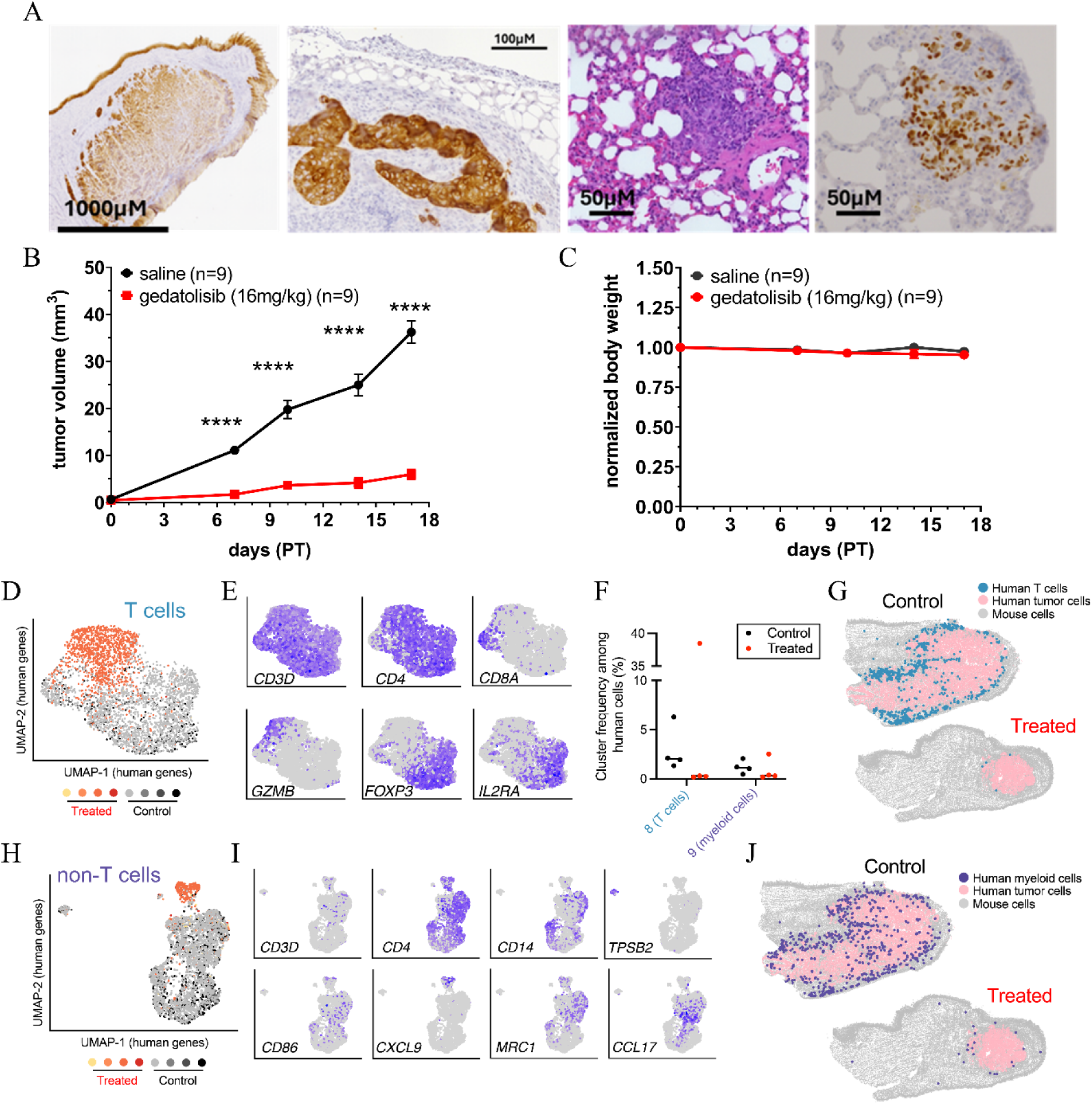
Gedatolisib suppresses tumor growth in humanized mice and shifts the TIME. **(A)** Cytokeratin staining of primary tumor (left most panel) and metastatic lymph node (2^nd^ left most panel) along with H&E (2^nd^ right most panel) and p40 (right most panel) of lung metastasis confirm an aggressive phenotype of HN30R8 in humanized mice. **(B)** *In vivo* data of orthotopic humanized model using HN30R8 cisplatin-resistant cell lines treated with 16 mg/kg of gedatolisib via tail vein (IV) administration every 3 days. The y-axis represents tumor volume (mm²), and the x-axis represents days post-injection following tumor visualization. Black data points indicate the control group, while red data points represent the gedatolisib-treated group. Data are presented as the mean ± standard error of the mean (SEM). P-values were calculated using a two-tailed Student’s t-test (* P ≤ 0.05, ** P ≤ 0.01, *** P ≤ 0.001, **** P ≤ 0.0001). **(C)** Normalized body weight representation of *in vivo* orthotopic xenograft models treated with 20mg/kg of gedatolisib. The y-axis represents body weight, and the x-axis denotes days post-injection following tumor visualization. Black data points indicate the control group, while red data points represent the gedatolisib-treated group. **(D)** Uniform manifold approximation and projection (UMAP) of tumor-infiltrating T cell gene expression as measured by the Xenium platform. **(E)** Expression of selected genes in tumor-infiltrating T cells. **(F)** Frequency of immune cell clusters among human cells within the tumor immune microenvironment (TIME). **(G)** Representative images showing the spatial distribution of T cells in control and treated tumors. **(H-I)** UMAP of non-T/myeloid immune cells in the tumor microenvironment and expression of selected genes. **(J)** Spatial distribution of myeloid cells within a treated and control tumor.

To study changes in the organization of the tumor immune microenvironment (TIME) following gedatolisib treatment, we employed the Xenium spatial transcriptomics platform to map gene expression at near single-cell resolution (**figure S15A-B)**. By employing hybridization probes against both human and mouse RNA targets, we could readily distinguish between human tumor and immune cells and surrounding mouse cells within the TME (**figure S15C-G**). We performed nearest-neighbor clustering and UMAP projection on gene expression signatures from single human cells within the TME, identifying four tumor cell clusters and two immune cell clusters (**figures 5D, 6A-B**). While CD4^+^ T cells predominated among T cells in both treated and control tumors, *FOXP3* expression was higher in T cells in control tumors, indicating sizeable regulatory T cell infiltration into the TME of untreated tumor cells. Gedatolisib treatment reduced the overall number of T cells, including *FOXP3*^+^ CD4^+^ T cells (**figure 5E-F**); these T cells were primarily located at the periphery of untreated tumors (**figure 5G**). Gedatolisib treatment also reduced the number of human myeloid cells within the TME (**figure 5G**). These cells were primarily CD4^+^ macrophages, with mixed expression of pro-inflammatory and immune suppressive genes such as *CD86*, *CXCL9*, *MRC1*, and *CCL17* (**figure 5H-I**). A small fraction of mast cells, identified by expression of *TPSB2,* were also detectable in control tumors (**figure 5I**). Spatially, myeloid cells predominantly localized to the tumor periphery, with minimal infiltration into the tumor parenchyma, but still infiltrated more extensively than T cells (**figure 5G, J**). Nearest-neighbor clustering identified four clusters of tumor cells with distinct patterns of gene expression (**figure 6A-C; supplementary tables 23-30**). Gene set enrichment analysis identified a cluster (cluster 1) defined by enrichment of hypoxia related genes and activation of the PI3K pathway; tumor clusters 2-4 were defined by differential expression of metabolic, cell adhesion, and cell cycle genes (**figure 6D; supplementary tables 27-30**). Gedatolisib treatment induced marked depletion of tumor cells from cluster 1 relative to control tumors (**figure 6E-F**). Consistent with their hypoxic signature, these cells were localized to the center of the tumor and could be demonstrated to be the furthermost from nearby endothelial cells (**figure 6G-I**). Loss of this cluster resulted in smaller tumors depleted of hypoxic regions, with more homogeneous vasculature. Global single-cell analysis of genes whose expression increased (>1.5 fold linear increase, adjusted p-value <0.05) as a function of treatment demonstrated enrichment of GO pathways involved in interferon and interleukin signaling, positive regulation of pyroptotic inflammatory response and positive regulation of MHC class I biosynthesis (**supplementary tables 23-30**). Conversely, down-regulated genes (>1.5 fold linear decrease, adjusted p-value <0.05) mapped to pathways involved in differentiation, morphogenesis and metabolism. Of note, critical metabolic genes including hexokinase 2, Glut3, enolase 2 and carbonic anhydrase 9 were noticeably downregulated following gedatolisib treatment (**supplementary tables 23-30**).

**Figure 6:**
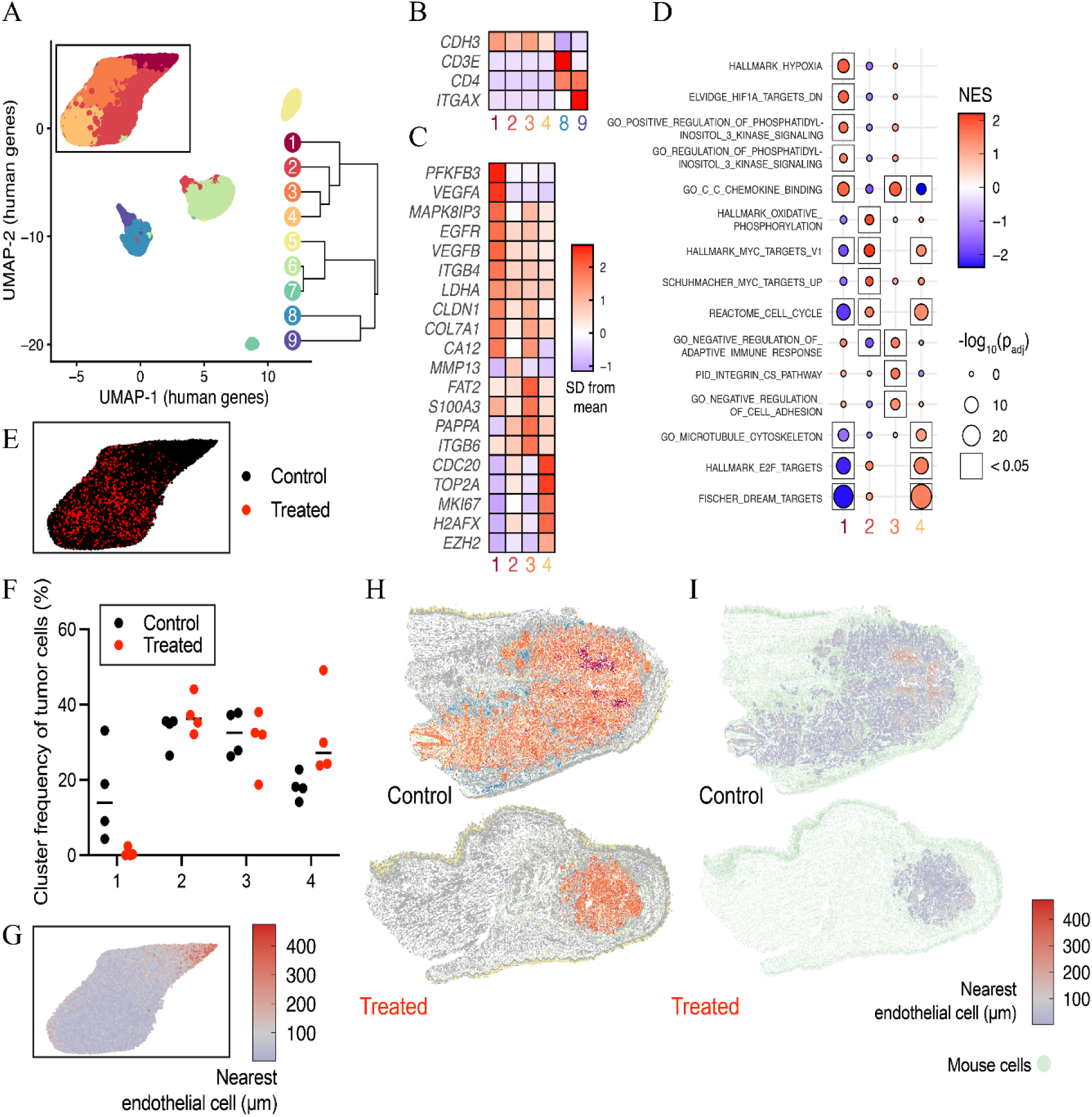
Gedatolisib treatment remodels tumor gene expression profiles and spatial organization. Four gedatolisib-treated and four control tumors were subjected to Xenium spatial transcriptomics analysis. **(A)** UMAP and nearest-neighbor clustering of human cells within the TME (n = 503,062 cells from eight mice). **(B)** Relative expression of lineage-defining genes reveals 4 tumor clusters (1–4) and two immune clusters (8 and 9) of T cells and myeloid cells, respectively. **(C)** Selected gene expression among tumor cell clusters. **(D)** Gene set enrichment analysis of gene expression within each tumor cell cluster compared to other tumor cells. **(E)** Inset of UMAP from panel (**A**) showing distribution of tumor cells from control or gedatolisib-treated ice. **(F)** Frequency of each tumor cluster in control and treated mice. **(G)** Inset of UMAP from panel **(A)** showing the distance of each cell from an endothelial cell. **(H)** Localization of human cell clusters within the TME in control and treated mice. Mouse cells are shown in gray. **(I)** Distance of human cells in the TME to the nearest endothelial cell; mouse cells are shown in green.

## Discussion

Effective targeting of specific signaling pathways has proven effective in cancer, as in the case of BRAF+MEK inhibition in *BRAF*-mutant melanoma and anaplastic thyroid cancer(31), EGFR inhibition in *EGFR*-mutant lung cancer and PI3Kinase inhibition in *PIK3CA*-mutant breast cancer (19,32–34) In the context of HNSCC, we have yet to define a targeted strategy with a sufficiently high therapeutic index that can achieve clinical relevance. In the current study, we chose the most aggressive phenotype of HNSCC that we can reproduce in preclinical models that match the human disease: chemo-radiation refractory disease, with associated suppressed immunity and a higher rate of regional and distant metastasis, all driven by hyperactivation of the Nrf2 pathway (10,11,20–23). Using a previous discovery that *NOTCH1* mutant cells demonstrated increased sensitivity to PI3K inhibition regardless of other genomic events (16,17), we sought to determine if this putative synergistic lethality can overcome the Nrf2-driven aggressive phenotype and result in meaningful anti-tumor activity for a new PI3Ki with a favorable toxicity profile. As shown here, classic PI3Kinase inhibitors like copanlisib, along with gedatolisib, demonstrate potent *in vitro* and *in vivo* antitumor activity in chemo-radiation-resistant HNSCC, albeit with a variable toxicity profile in favor of gedatolisib. Suppression of PI3Kinase signaling is both profound and durable at low nanomolar concentrations which speaks well for clinical translation. Surprisingly, gedatolisib demonstrated significant activity against HNSCC cell lines irrespective of HPV-association and importantly, even in *NOTCH1* wild-type cell lines, which is encouraging from the perspective of clinical translation. Overall, the data point toward a complex metabolic and pathway-driven shutdown of cell proliferation with the induction of cell arrest initially and with the induction of autophagy, senescence, and cell death at later stages. Although metabolic changes appear to be of a secondary order, the convergence of metabolic disruption of carnitine derivatives points to altered fatty acid metabolism as a potential target for further enhancement of gedatolisib effectiveness.

We used four distinct preclinical models to test the potential for gedatolisib, in part because of the previously disappointing translation of other PI3Kinase inhibitors (35–37). In the conventional orthotopic model, the drug demonstrated potent antitumor activity, similar to what we have described for other effective agents over the last 10 years, with a tolerable toxicity profile. Our preliminary analysis of efficacy in the lung metastasis setting, which is the most common site for distant metastasis in HNSCC, and the measured effectiveness in the presence of a (partially) functional immune system in the humanized mouse model is supportive of gedatolisib effectiveness as a single agent in otherwise treatment-refractory disease. Although targeted agents can be used alone in other solid tumors, in HNSCC that is generally not the case. Nevertheless, in the context of cisplatin and radiation refractory disease, which generally has an adverse anti-tumor immune profile, all driven by Nrf2 hyperactivation, we thought that a single agent approach would be most consistent with the traditional preclinical modeling required for phase I/II trials in the recurrent/metastatic disease setting (37). This is particularly relevant since failure of immune checkpoint inhibitors, which we have now linked to Nrf2 hyperactivation in preclinical murine models (12), essentially mandates subsequent on-trial treatment of HNSCC. The potential for combinatorial strategies is supported by our data, as gedatolisib does not appear to generate substantial systemic immune suppression at effective doses, which would negatively overlap with the systemic effects of cisplatin in treatment-naïve disease. Reduced levels of members of the *AKR* family speak to potential direct re-sensitization to both cisplatin and radiation (19,38). Although the spatial transcriptomic data demonstrate a profound reduction in tumor cell clusters demonstrating activation of hypoxia and altered metabolism, which are encouraging from the perspective of restoring chemo-radiation effectiveness, we caution that a reduction in tumor volume can at least partially explain these effects. Similarly, at least a portion of the shifts in anti-tumor immunity may likely be driven by decreased tumor size. Nevertheless, smaller tumors, void of hypoxic regions and inhibitory Tregs, in which Nrf2 targets are down-regulated do present a very attractive opportunity for additional combinatorial work. Finally, the disruptions measured in basic metabolic pathways both *in vitro* and *in vivo* present additional opportunities for combinatorial testing.

Future planned studies will determine whether gedatolisib effectiveness as a single drug or as a chemosensitizing agent is maintained across variable *PIK3CA* and *NOTCH1* mutational backgrounds in HNSCC and whether gedatolisib can restore the activity of immune checkpoint inhibitors in syngeneic and genetically engineered murine models of Nrf2 hyperactivation. Since Nrf2 activation can be detected in the pre-treatment setting, as we and others have shown using bulk RNA-seq and more targeted platforms, (10) a potential biomarker-informed strategy could be leveraged to maximize the therapeutic index of gedatolisib and other compounds in this class.

## Data availability

The data generated in this study are available in the article and its supplementary files.

## Funding to acknowledge

This work was supported by the Sid Richardson Foundation, by the National Cancer Institute through U54CA274321 and R01CA280980, by the Veterans Affairs Basic Science Research and Development division (1I01BX006380), and by the National Institutes of Health (1R01CA235620 to FMJ, MJF). N.P. is supported by the CPRIT Proteomics and Metabolomics Core Facility (RP210227), NIH (P30 CA125123, R01CA282282, P42ES027725), S10OD032218 and Dan L. Duncan Cancer Center. CC is partially supported by CPRIT RP210227 and RP200504, NIH/NCI P30 shared resource grant CA125123, NIH/NIEHS P42 ES027725 and P30 ES030285. W.H.H. is supported by NIH (R00AI153736 and R37CA285289) and a first-time tenure-track award (RR220069) from the Cancer Prevention and Research Institute of Texas (CPRIT). W.H.H. is a CPRIT Scholar in Cancer Research. This manuscript does not represent the views and opinions of the US Department of Veterans Affairs or the National Institutes of Health.

## Reference

1. Sung H, Ferlay J, Siegel RL, Laversanne M, Soerjomataram I, Jemal A, et al. Global Cancer Statistics 2020: GLOBOCAN Estimates of Incidence and Mortality Worldwide for 36 Cancers in 185 Countries. CA Cancer J Clin 2021;71(3):209–49 doi 10.3322/caac.21660.

2. Harrington KJ, Burtness B, Greil R, Soulieres D, Tahara M, de Castro G, Jr., et al. Pembrolizumab With or Without Chemotherapy in Recurrent or Metastatic Head and Neck Squamous Cell Carcinoma: Updated Results of the Phase III KEYNOTE-048 Study. J Clin Oncol 2023;41(4):790–802 doi 10.1200/JCO.21.02508.

3. Agrawal N, Frederick MJ, Pickering CR, Bettegowda C, Chang K, Li RJ, et al. Exome sequencing of head and neck squamous cell carcinoma reveals inactivating mutations in NOTCH1. Science 2011;333(6046):1154–7 doi 10.1126/science.1206923.

4. Cancer Genome Atlas N. Comprehensive genomic characterization of head and neck squamous cell carcinomas. Nature 2015;517(7536):576–82 doi 10.1038/nature14129.

5. Pickering CR, Zhang J, Yoo SY, Bengtsson L, Moorthy S, Neskey DM, et al. Integrative genomic characterization of oral squamous cell carcinoma identifies frequent somatic drivers. Cancer Discov 2013;3(7):770–81 doi 10.1158/2159-8290.CD-12-0537.

6. Stransky N, Egloff AM, Tward AD, Kostic AD, Cibulskis K, Sivachenko A, et al. The mutational landscape of head and neck squamous cell carcinoma. Science 2011;333(6046):1157–60 doi 10.1126/science.1208130.

7. Ferris RL, Blumenschein G, Jr., Fayette J, Guigay J, Colevas AD, Licitra L, et al. Nivolumab for Recurrent Squamous-Cell Carcinoma of the Head and Neck. N Engl J Med 2016;375(19):1856–67 doi 10.1056/NEJMoa1602252.

8. Taberna M, Oliva M, Mesia R. Cetuximab-Containing Combinations in Locally Advanced and Recurrent or Metastatic Head and Neck Squamous Cell Carcinoma. Front Oncol 2019;9:383 doi 10.3389/fonc.2019.00383.

9. Fu R, Zhao B, Chen M, Fu X, Zhang Q, Cui Y, et al. Moving beyond cisplatin resistance: mechanisms, challenges, and prospects for overcoming recurrence in clinical cancer therapy. Med Oncol 2023;41(1):9 doi 10.1007/s12032-023-02237-w.

10. Osman AA, Arslan E, Bartels M, Michikawa C, Lindemann A, Tomczak K, et al. Dysregulation and Epigenetic Reprogramming of NRF2 Signaling Axis Promote Acquisition of Cisplatin Resistance and Metastasis in Head and Neck Squamous Cell Carcinoma. Clin Cancer Res 2023;29(7):1344–59 doi 10.1158/1078-0432.CCR-22-2747.

11. Yu W, Chen Y, Putluri N, Osman A, Coarfa C, Putluri V, et al. Evolution of cisplatin resistance through coordinated metabolic reprogramming of the cellular reductive state. Br J Cancer 2023;128(11):2013–24 doi 10.1038/s41416-023-02253-7.

12. Ahmed KM, Veeramachaneni R, Deng D, Putluri N, Putluri V, Cardenas MF, et al. Glutathione peroxidase 2 is a metabolic driver of the tumor immune microenvironment and immune checkpoint inhibitor response. J Immunother Cancer 2022;10(8) doi 10.1136/jitc-2022-004752.

13. Liu GH, Chen T, Zhang X, Ma XL, Shi HS. Small molecule inhibitors targeting the cancers. MedComm (2020) 2022;3(4):e181 doi 10.1002/mco2.181.

14. Janku F, Yap TA, Meric-Bernstam F. Targeting the PI3K pathway in cancer: are we making headway? Nat Rev Clin Oncol 2018;15(5):273–91 doi 10.1038/nrclinonc.2018.28.

15. Mazumdar T, Byers LA, Ng PK, Mills GB, Peng S, Diao L, et al. A comprehensive evaluation of biomarkers predictive of response to PI3K inhibitors and of resistance mechanisms in head and neck squamous cell carcinoma. Mol Cancer Ther 2014;13(11):2738–50 doi 10.1158/1535-7163.MCT-13-1090.

16. Shah PA, Huang C, Li Q, Kazi SA, Byers LA, Wang J, et al. NOTCH1 Signaling in Head and Neck Squamous Cell Carcinoma. Cells 2020;9(12) doi 10.3390/cells9122677.

17. Sambandam V, Frederick MJ, Shen L, Tong P, Rao X, Peng S, et al. PDK1 Mediates NOTCH1-Mutated Head and Neck Squamous Carcinoma Vulnerability to Therapeutic PI3K/mTOR Inhibition. Clin Cancer Res 2019;25(11):3329–40 doi 10.1158/1078-0432.CCR-18-3276.

18. Johnson FM, Janku F, Gouda MA, Tran HT, Kawedia JD, Schmitz D, et al. Inhibition of the Phosphatidylinositol-3 Kinase Pathway Using Bimiralisib in Loss-of-Function NOTCH1-Mutant Head and Neck Cancer. Oncologist 2022;27(12):1004–e926 doi 10.1093/oncolo/oyac185.

19. Curigliano G, Shapiro GI, Kristeleit RS, Abdul Razak AR, Leong S, Alsina M, et al. A Phase 1B open-label study of gedatolisib (PF-05212384) in combination with other anti-tumour agents for patients with advanced solid tumours and triple-negative breast cancer. Br J Cancer 2023;128(1):30–41 doi 10.1038/s41416-022-02025-9.

20. Yassin-Kassab A, Chatterjee S, Khan N, Wang N, Sandulache VC, Huang EH, et al. p90RSK pathway inhibition synergizes with cisplatin in TMEM16A overexpressing head and neck cancer. BMC Cancer 2024;24(1):233 doi 10.1186/s12885-024-11892-9.

21. Pifer PM, Yang L, Kumar M, Xie T, Frederick M, Hefner A, et al. FAK drives resistance to therapy in HPV-negative head and neck cancer in a p53-dependent manner. Clin Cancer Res 2023 doi 10.1158/1078-0432.CCR-23-0964.

22. Yu W, Chen Y, Putluri N, Coarfa C, Robertson MJ, Putluri V, et al. Acquisition of Cisplatin Resistance Shifts Head and Neck Squamous Cell Carcinoma Metabolism toward Neutralization of Oxidative Stress. Cancers (Basel) 2020;12(6) doi 10.3390/cancers12061670.

23. Pham D, Deter CJ, Reinard MC, Gibson GA, Kiselyov K, Yu W, et al. Using Ligand-Accelerated Catalysis to Repurpose Fluorogenic Reactions for Platinum or Copper. ACS Cent Sci 2020;6(10):1772–88 doi 10.1021/acscentsci.0c00676.

24. Frederick M, Skinner HD, Kazi SA, Sikora AG, Sandulache VC. High expression of oxidative phosphorylation genes predicts improved survival in squamous cell carcinomas of the head and neck and lung. Sci Rep 2020;10(1):6380 doi 10.1038/s41598-020-63448-z.

25. Hao Y, Stuart T, Kowalski MH, Choudhary S, Hoffman P, Hartman A, et al. Dictionary learning for integrative, multimodal and scalable single-cell analysis. Nat Biotechnol 2024;42(2):293–304 doi 10.1038/s41587-023-01767-y.

26. Hadley W. Ggplot2. New York, NY: Springer Science+Business Media, LLC; 2016. pages cm p.

27. McInnes L, Healy J, Melville J. UMAP: Uniform Manifold Approximation and Projection for Dimension Reduction. arXiv 2020;1802.03426.

28. Korotkevich G, Sukhov V, Budin N, Shpak B, Artyomov MN, Sergushichev A. Fast gene set enrichment analysis. bioRvix 2021 doi 10.1101/060012.

29. Chang WM, Chang YC, Yang YC, Lin SK, Chang PM, Hsiao M. AKR1C1 controls cisplatin-resistance in head and neck squamous cell carcinoma through cross-talk with the STAT1/3 signaling pathway. J Exp Clin Cancer Res 2019;38(1):245 doi 10.1186/s13046-019-1256-2.

30. Herrero-Sanchez MC, Rodriguez-Serrano C, Almeida J, San Segundo L, Inoges S, Santos-Briz A, et al. Targeting of PI3K/AKT/mTOR pathway to inhibit T cell activation and prevent graft-versus-host disease development. J Hematol Oncol 2016;9(1):113 doi 10.1186/s13045-016-0343-5.

31. Maniakas A, Dadu R, Busaidy NL, Wang JR, Ferrarotto R, Lu C, et al. Evaluation of Overall Survival in Patients With Anaplastic Thyroid Carcinoma, 2000-2019. JAMA Oncol 2020;6(9):1397–404 doi 10.1001/jamaoncol.2020.3362.

32. Andre F, Ciruelos E, Rubovszky G, Campone M, Loibl S, Rugo HS, et al. Alpelisib for PIK3CA-Mutated, Hormone Receptor-Positive Advanced Breast Cancer. N Engl J Med 2019;380(20):1929–40 doi 10.1056/NEJMoa1813904.

33. Sweetlove M, Wrightson E, Kolekar S, Rewcastle GW, Baguley BC, Shepherd PR, et al. Inhibitors of pan-PI3K Signaling Synergize with BRAF or MEK Inhibitors to Prevent BRAF-Mutant Melanoma Cell Growth. Front Oncol 2015;5:135 doi 10.3389/fonc.2015.00135.

34. Lu S, Kato T, Dong X, Ahn MJ, Quang LV, Soparattanapaisarn N, et al. Osimertinib after Chemoradiotherapy in Stage III EGFR-Mutated NSCLC. N Engl J Med 2024;391(7):585–97 doi 10.1056/NEJMoa2402614.

35. Fukuhara N, Maruyama D, Hatake K, Nagai H, Makita S, Kamezaki K, et al. Safety and antitumor activity of copanlisib in Japanese patients with relapsed/refractory indolent non-Hodgkin lymphoma: a phase Ib/II study. Int J Hematol 2023;117(1):100–9 doi 10.1007/s12185-022-03455-0.

36. Damodaran S, Zhao F, Deming DA, Mitchell EP, Wright JJ, Gray RJ, et al. Phase II Study of Copanlisib in Patients With Tumors With PIK3CA Mutations: Results From the NCI-MATCH ECOG-ACRIN Trial (EAY131) Subprotocol Z1F. J Clin Oncol 2022;40(14):1552–61 doi 10.1200/JCO.21.01648.

37. Marret G, Isambert N, Rezai K, Gal J, Saada-Bouzid E, Rolland F, et al. Phase I trial of copanlisib, a selective PI3K inhibitor, in combination with cetuximab in patients with recurrent and/or metastatic head and neck squamous cell carcinoma. Invest New Drugs 2021;39(6):1641–8 doi 10.1007/s10637-021-01152-z.

38. Colombo I, Genta S, Martorana F, Guidi M, Frattini M, Samartzis EP, et al. Phase I Dose-Escalation Study of the Dual PI3K-mTORC1/2 Inhibitor Gedatolisib in Combination with Paclitaxel and Carboplatin in Patients with Advanced Solid Tumors. Clin Cancer Res 2021;27(18):5012–9 doi 10.1158/1078-0432.CCR-21-1402.

