## Supplemental Figures for "Bypassing cisplatin resistance in Nrf2 hyperactivated head and neck cancer through effective PI3Kinase targeting"

#### SUPPLEMENTAL FIGURES AND LEGENDS

Figure S1

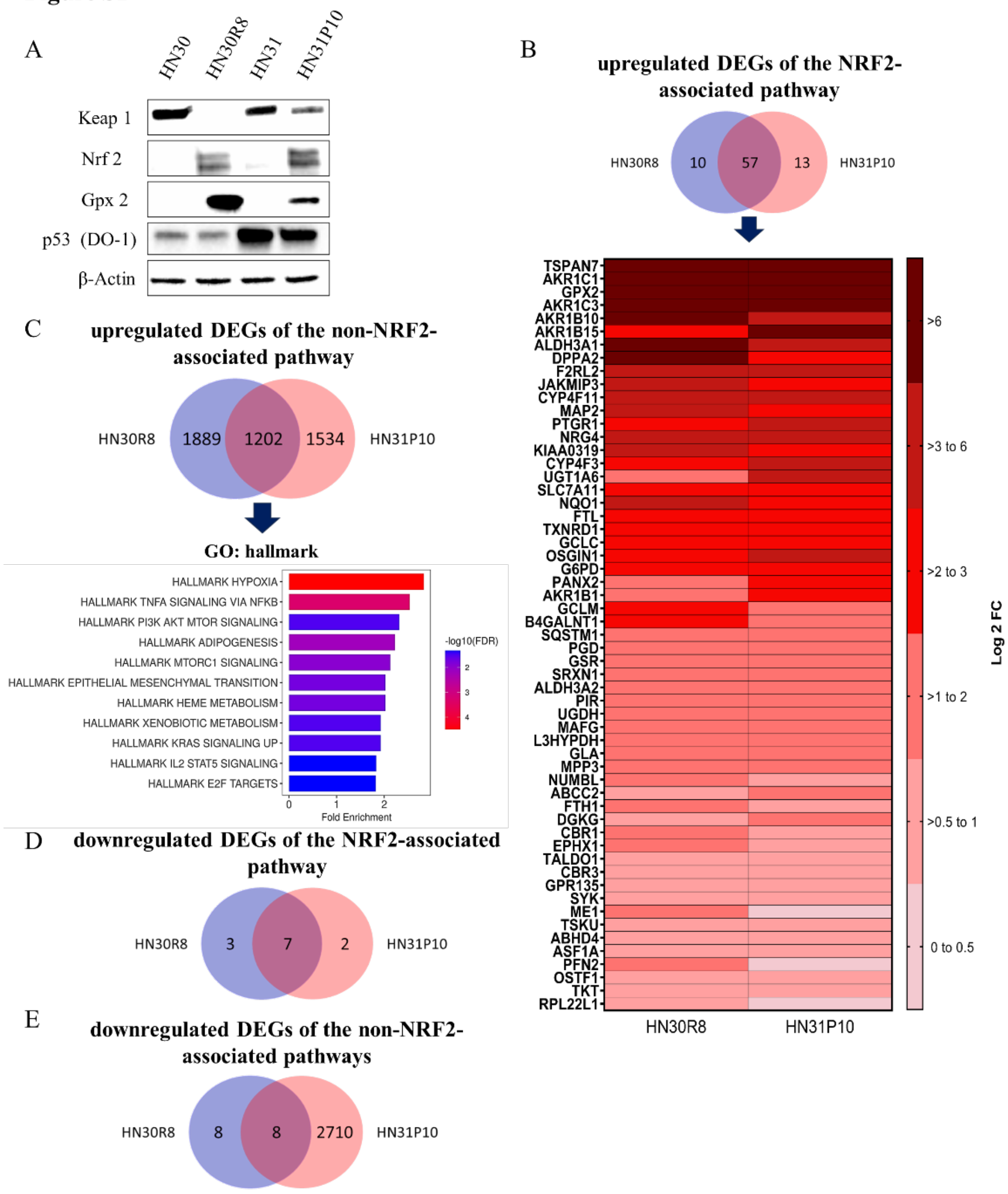


**Figure S1. PI3K pathway activity in cisplatin-resistant HNSCC cell lines. (A)** Western blot panel showing the basal expression of target proteins in HN30, HN31 and their cisplatin-resistant counterparts. β-actin serves as loading control. **(B)** Heatmap illustrates upregulated differentially expressed genes (DEGs) of the nrf2-asssociated pathway in HN30R8 and HN31P10 cisplatin-resistant cell lines compared to their parental lines, derived from a Venn diagram using in vitro RNA-seq data. Gene expression is represented as log2 fold change, with red indicating upregulated genes. **(C)** upregulated DEGs of the non-nrf2-asssociated pathway analysis of HN30R8 and HN31P10 cisplatin-resistant cell lines, showing Hallmark gene enrichment for upregulated genes, presented as a bar graph and sorted by log10 False Discovery Rate (FDR) with a cutoff ≤0.05 and fold enrichment. **(D)** Venn diagram analysis of downregulated DEGs of the Nrf2-associated pathway between HN30R8 and HN31P10. **(E)** Venn diagram analysis downregulated DEGs of the non-Nrf2-associated pathway between HN30R8 and HN31P10.

**Figure S2**

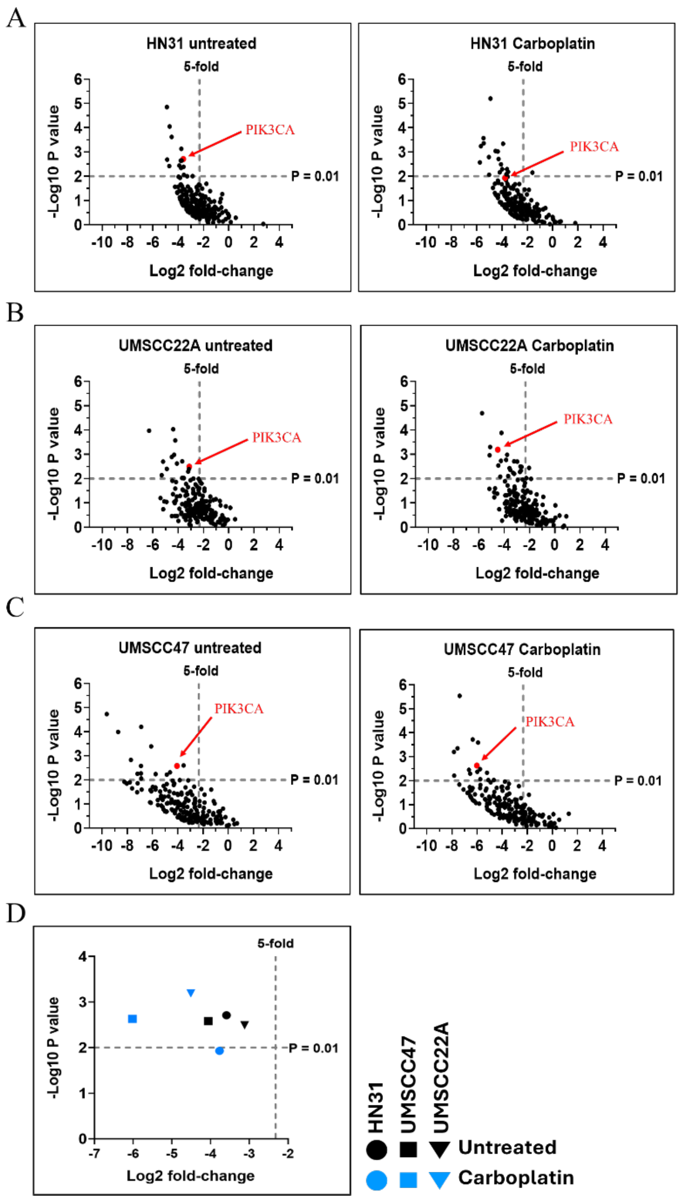


**Figure S2. NOTCH1 mutant HNSCC cell lines depend on PIK3CA for baseline cell fitness and after genotoxic treatments. (A,B)** HN31, **(C,D)** UMSCC22A, or **(E, F)** UMSCC47 were screened with an shRNA library targeting 195 genes, including PIK3CA (red dots) and the RSA method was used to calculate log2 fold changes and P-values for each gene based on dropout compared to tumors grown in mice receiving no treatment **(A, C , E)**, carboplatin **(B, D, F)**. **(G)** P-values and log2 fold changes only for *PIK3CA* were compared across cell lines and treatments.

**Figure S3**

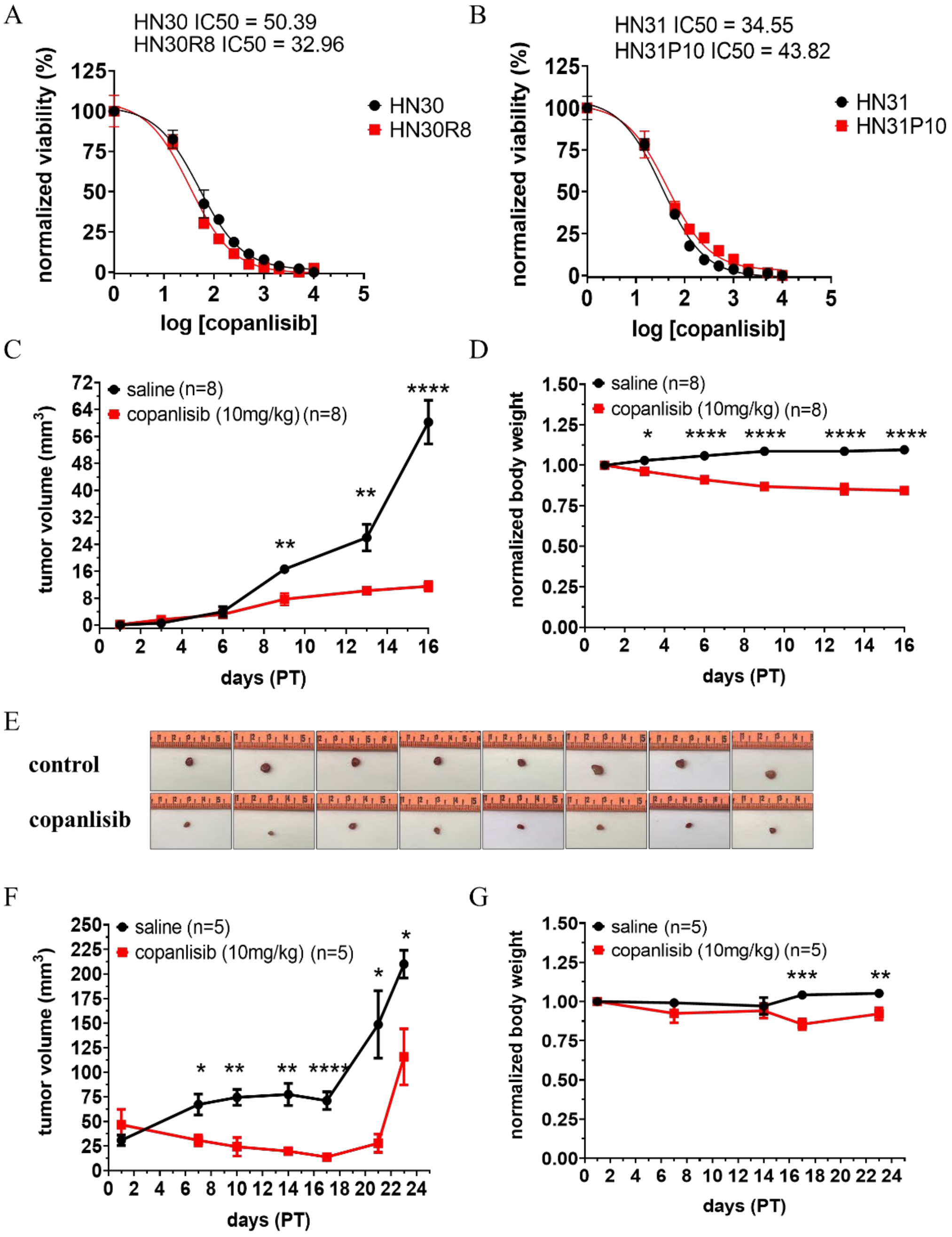


**Figure S3. Copanlisib reduces HNSCC cisplatin-resistant tumor growth accompanied by induction of adverse effect on body weight. (A-B)** HNSCC cell lines were treated with copanlisib for 72 hours and assessed using a Resazurin assay. The sigmoid graph plots normalized cell viability (y-axis) against the logarithm of copanlisib concentration (x-axis) to determine the $IC_{50}$. Black represents parental cell lines, while red represents cisplatin-resistant cell lines. Data are presented as the mean ± standard error of the mean (SEM). P-values were calculated using a two-tailed Student's t-test (* P ≤ 0.05, ** P ≤ 0.01, *** P ≤

0.001, **** P ≤ 0.0001). **(C)** In vivo data from orthotopic xenograft model using HN30R8 cisplatin-resistant cell lines treated with 10 mg/mg of copanlisib via intraperitoneal (IP) administration every 2 days. The y-axis represents tumor volume ($mm^2$), and the x-axis represents days post-injection following tumor visualization. Black data points indicate the control group, while red data points represent the copanlisib-treated group. **(D)** Normalized body weight representation of *in vivo* orthotopic xenograft model treated with 10mg/kg of copanlisib. The y-axis represents body weight (grams), and the x-axis denotes days post-injection following tumor visualization. Black data points indicate the control group, while red data points represent the copanlisib-treated group. **(E)** Gross images of extracted tumor from orthotopic model treated with copanlisib. **(F*)*** *In vivo* data from subcutaneous xenograft model using HN30R8 cisplatin-resistant cell lines, treated with 10 mg/kg of copanlisib via intraperitoneal (IP) administration every 2 days. Y-axis represents body weight (grams), and the x-axis denotes days post-injection following tumor visualization. Black data points indicate the control group, while red data points represent the copanlisib-treated group. **(G)** Normalized body weight representation of in vivo subcutaneous xenograft model treated with 10mg/kg of copanlisib. The y-axis represents body weight (grams), and the x-axis denotes days post-injection following tumor visualization. Black data points indicate the control group, while red data points represent the copanlisib-treated group. Data are presented as the mean ± standard error of the mean (SEM). P-values were determined using a two-tailed Student's t-test (ns, not significant; * P ≤ 0.05). D. The images of tumor extracted from orthotopic mice.

**Figure S4**

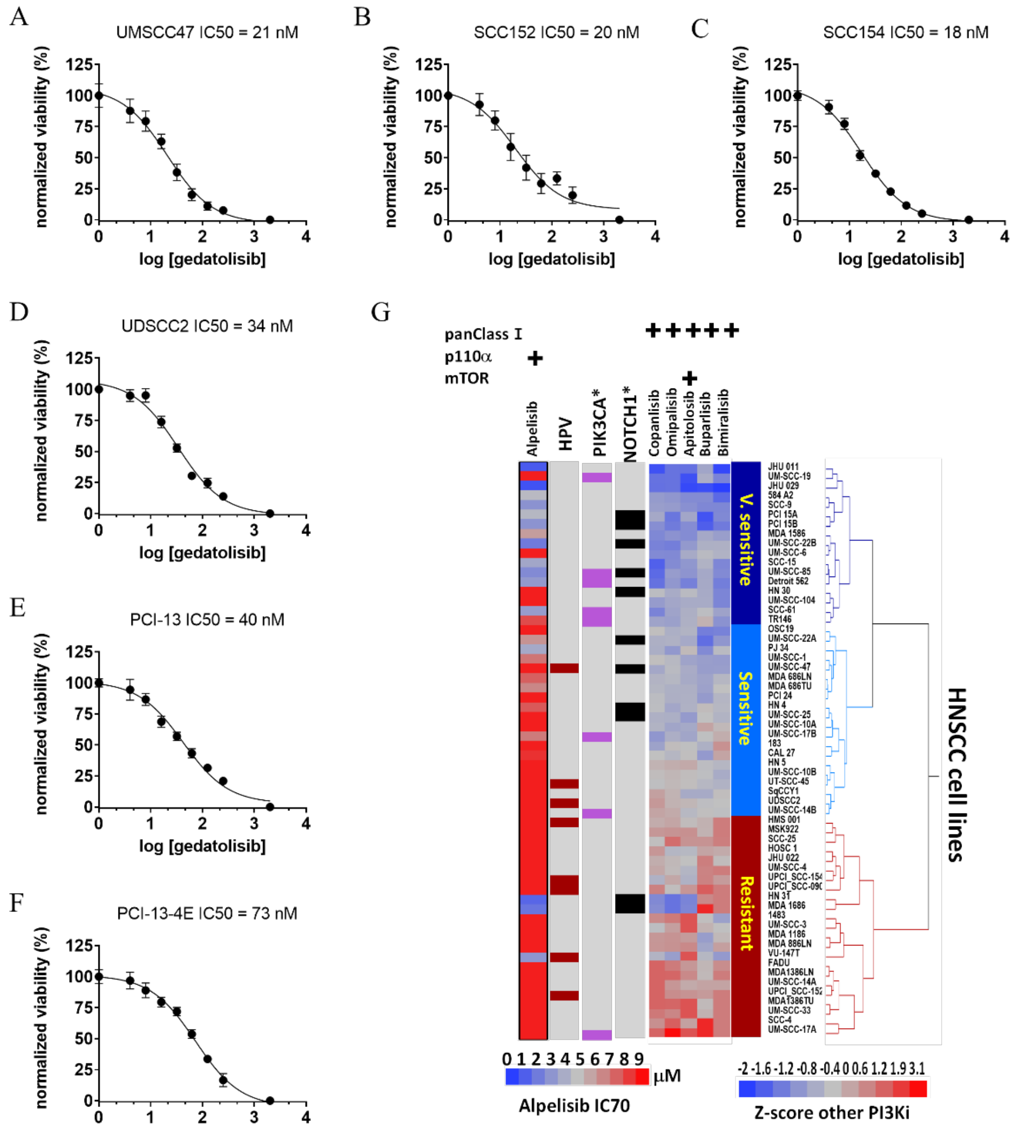


| | NOTCH1 mut | PI3K mut | Wt | HPV+ |
|---|---|---|---|---|
| V. sensitive | 5 | 5 | 8 | 0 |
| Sensitive | 4 | 2 | 14 | 3 |
| Resistant | 2 | 1 | 20 | 5 |

**Figure S4. Gedatolisib drug sensitivity in various HNSCC cell lines. (A-F)** HPV-associated (UMSCC47, UDSCC2, SCC152, SCC154) and HPV-independent (PCI13) HNSCC lines as well as an additional cisplatin-resistant (PCI134E) line were treated with gedatolisib for 72 hours and assessed using a Resazurin assay. The sigmoid graph plots normalized cell viability (y-axis) against the logarithm of gedatolisib concentration (x-axis) to determine the $IC_{50}$. **(G)** Differential analysis of PI3Ki sensitivity in a large panel of HNSCC cell lines as a function of HPV status and *NOTCH1* and *PIK3CA* mutational (mut) status. The data are presented in z-score scale and cells were clustered into three groups: resistant, sensitive, and very sensitive (v.sensitive).

**Figure S5**

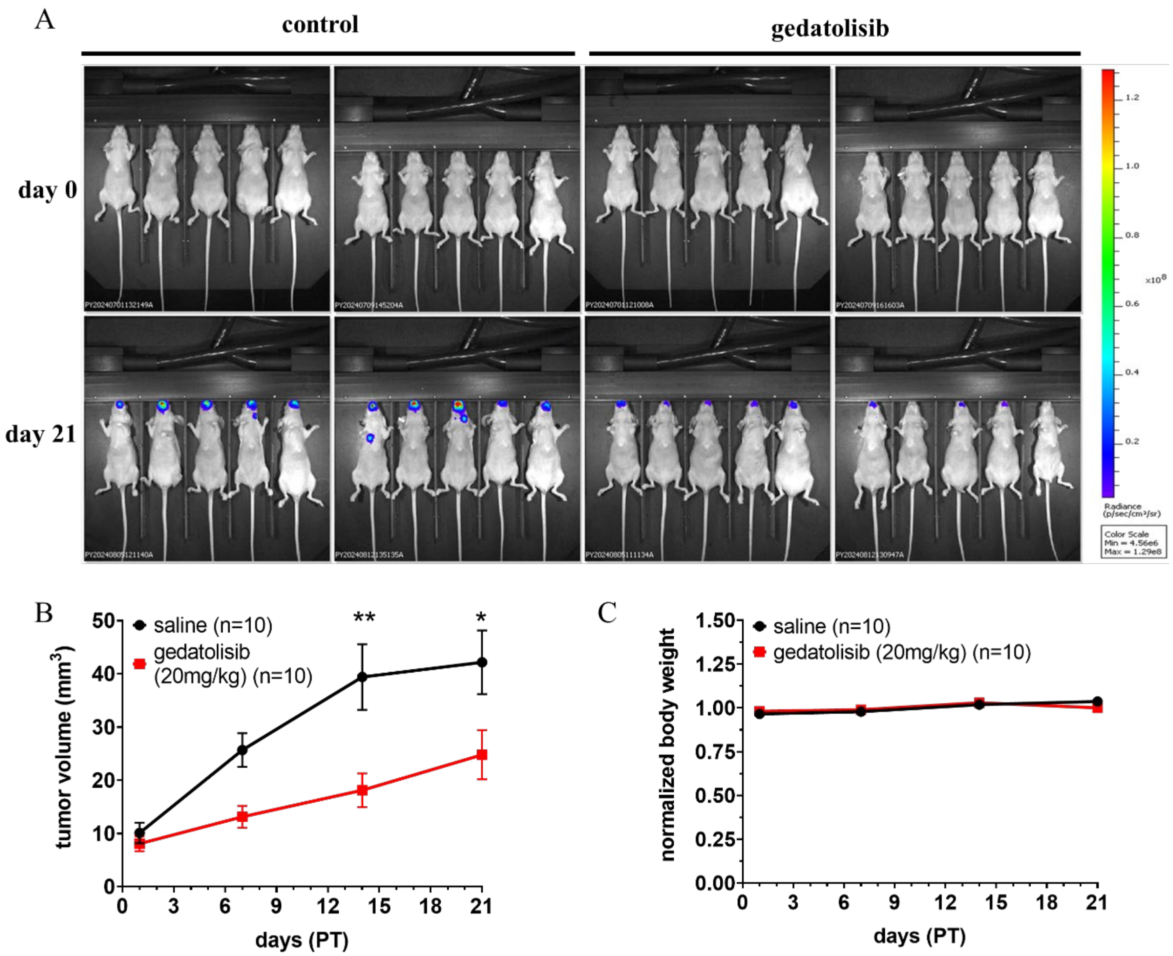


**Figure S5. Gedatolisib suppresses cisplatin-resistant HNSCC tumor growth. (A)** IVIS signal shows luciferase activity in mice bearing HN30R8 tumors, at day 0 and day 21 post-treatment. **(B)** In vivo data from orthotopic xenograft models using HN30R8 cisplatin-resistant cell lines treated with 20 mg/kg of gedatolisib via tail vein (IV) administration every 3 days. The y-axis represents tumor volume ($mm^3$), and the x-axis represents days post-injection following tumor visualization. Black data points indicate the control group, while red data points represent the gedatolisib-treated group. Data are presented as the mean ± standard error of the mean (SEM). P-values were calculated using a two-tailed Student's t-test (* $P \leq 0.05$, ** $P \leq 0.01$, *** $P \leq 0.001$, **** $P \leq 0.0001$). **(C)** Normalized body weight representation of in vivo orthotopic xenograft models treated with 20mg/kg of gedatolisib. Y-axis represents body weight (grams), and the x-axis denotes days post-injection following tumor visualization. Black data points indicate the control group, while red data points represent the gedatolisib-treated group.

#### Figure S6

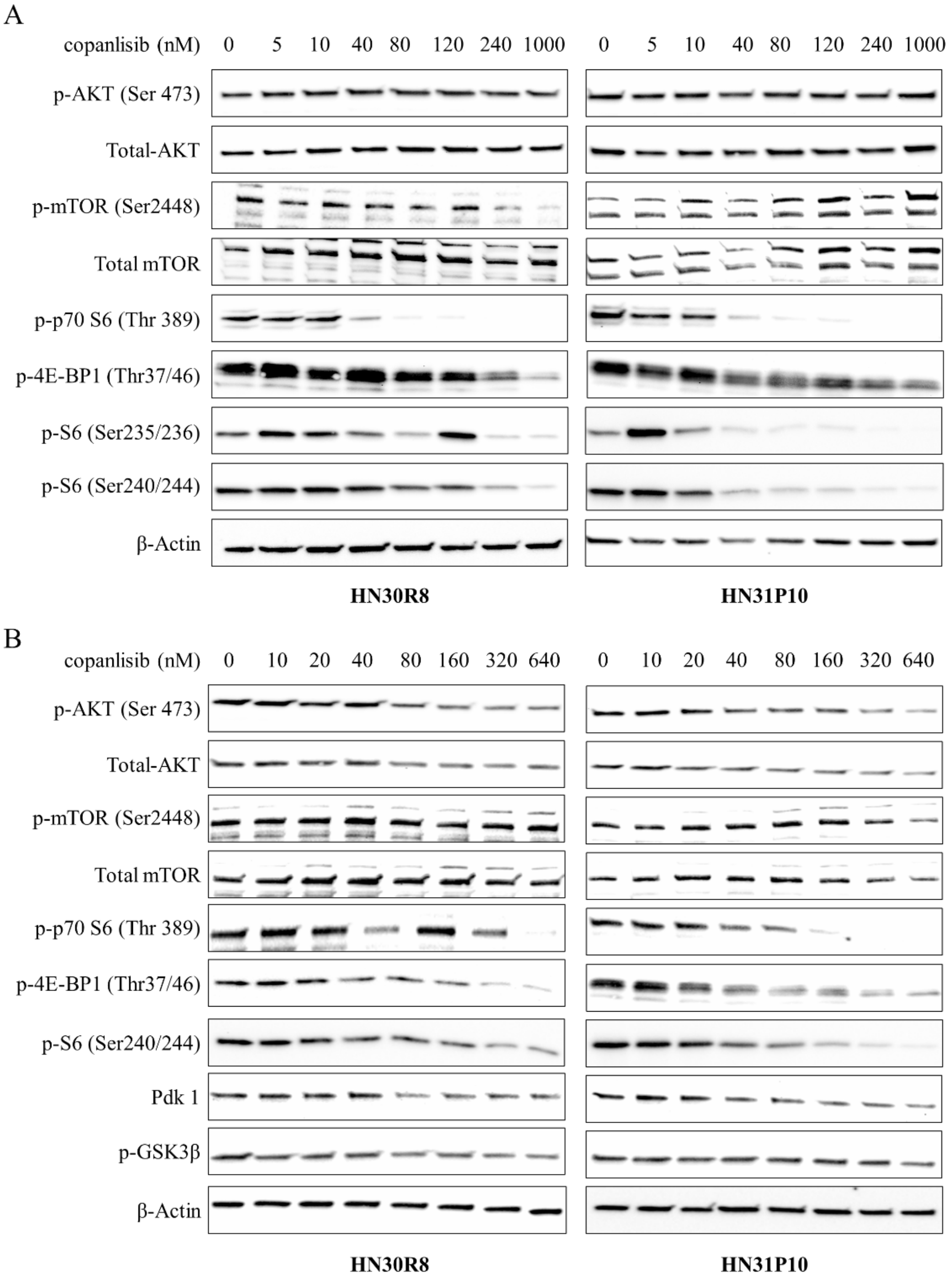


**Figure S6. Copanlisib effects on PI3K targets. (A)** Representative Western blot data showing selected PI3K downstream target proteins. HNSC cisplatin-resistant cell lines, HN30R8 (left panel) and HN31P10 (right panel), were treated with copanlisib and/or control for 6 hours at the indicated drug concentrations (nM). β-Actin was used as a loading control. **(B)** Representative Western blot data showing selected PI3K downstream target proteins. HNSC cisplatin-resistant cell lines, HN30R8 (left panel) and HN31P10 (right panel), were treated with copanlisib and/or control for 48 hours at the indicated drug concentrations (nM). β-Actin was used as a loading control.

**Figure S7**

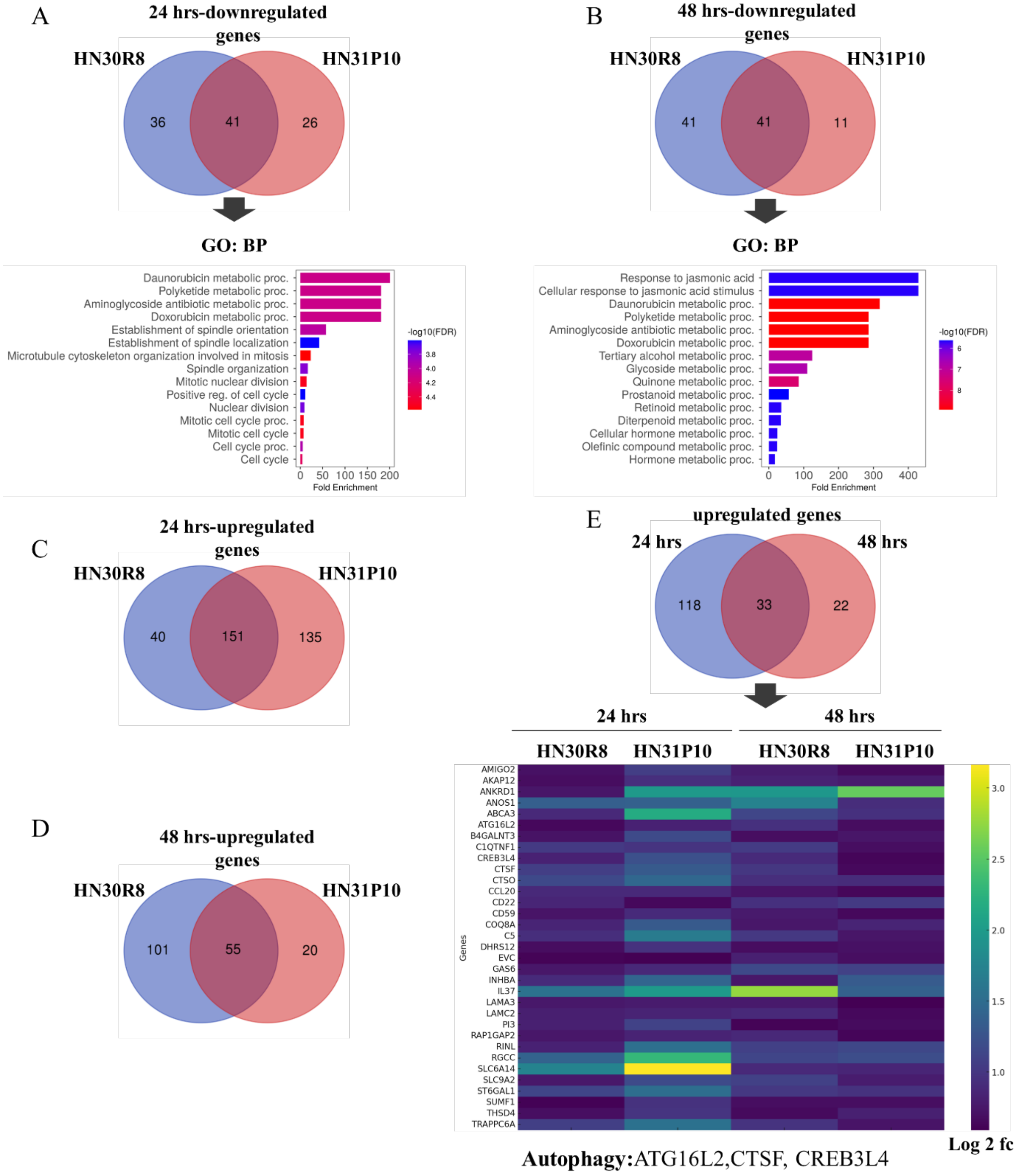


**Figure S7. Transcriptomic profile of gedatolisib. (A)** Differentially expressed genes (DEGs) analysis shows common downregulated genes in the cisplatin-resistant HN30R8 and HN31P10 cell lines after 24 hours of gedatolisib treatment, as illustrated by the Venn diagram. Bar graphs represent Gene Ontology (GO) analysis of biological processes (BP), with the y-axis depicting enriched pathways and the x-axis showing fold enrichment. Data were filtered based on a false discovery rate (FDR) threshold of 0.05, with significance criteria set at a fold change greater than 1.5. **(B)** DEG analysis displays downregulated overlapping genes in the cisplatin-resistant HN30R8 and HN31P10 cell lines after 48 hours of gedatolisib exposure, as shown in the Venn diagram. Bar graphs represent Gene Ontology (GO) analysis of biological processes (BP), with the y-axis depicting enriched pathways and the x-axis showing fold enrichment. Data were filtered based on a false discovery rate (FDR) threshold of 0.05, with significance criteria set at a fold change greater than 1.5. **(C)** The Venn diagram illustrates DEG analysis of upregulated overlapping genes in the cisplatin-resistant HN30R8 and HN31P10 cell lines after 24 hours of gedatolisib treatment. **(D)** DEG analysis illustrates upregulated overlapping genes in the cisplatin-resistant HN30R8 and HN31P10 cell lines after 48 hours of gedatolisib exposure, as shown in the Venn diagram. **(E)** The analysis compares common upregulated genes in HN30R8 and HN31P10 cisplatin-resistant cell lines at both 24-hour and 48-hour timepoints, with the Venn diagram highlighting the overlapping genes and shared upregulation between the two timepoints. A heatmap provides a visual representation of the differential expression of these common genes across both exposure periods, with color bar indicating the log2 fold change (FC).

**Figure S8**

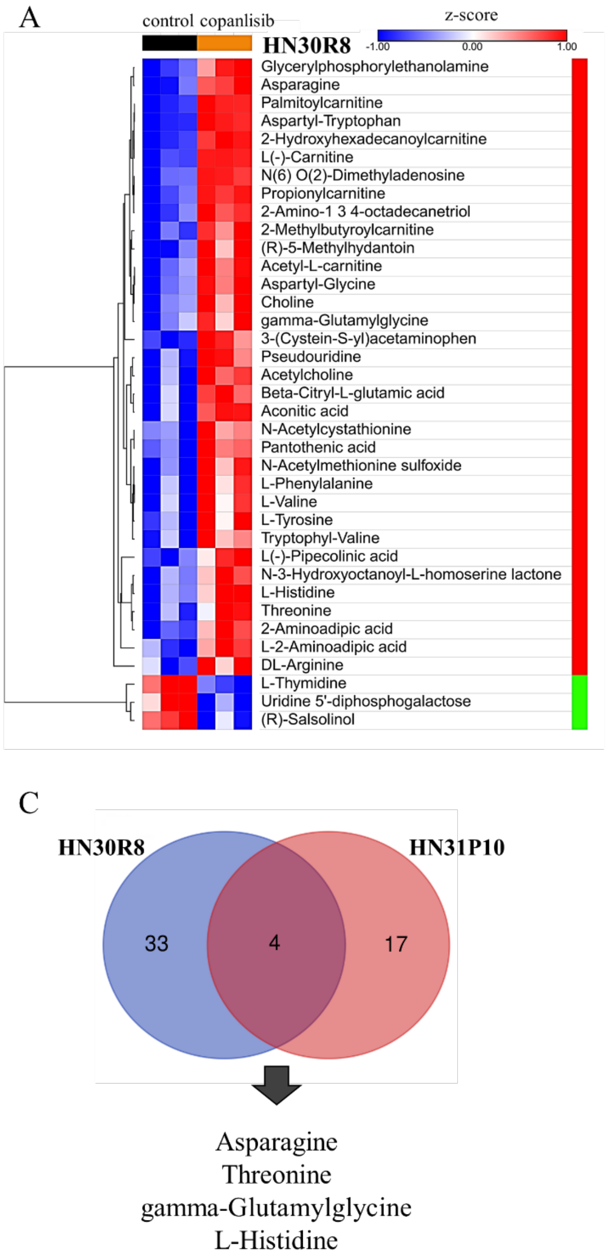


**Figure S8. Limited copanlisib influence on steady state metabolism in HNSCC cell lines. (A-C)** Heatmap of differentially-abundant metabolites in HN30R8 and HN31P10 cells treated with vehicle (control) or copanlisib for 24 hours, along with a Venn diagram highlighting the common metabolites in both cell lines.

**Figure S9**

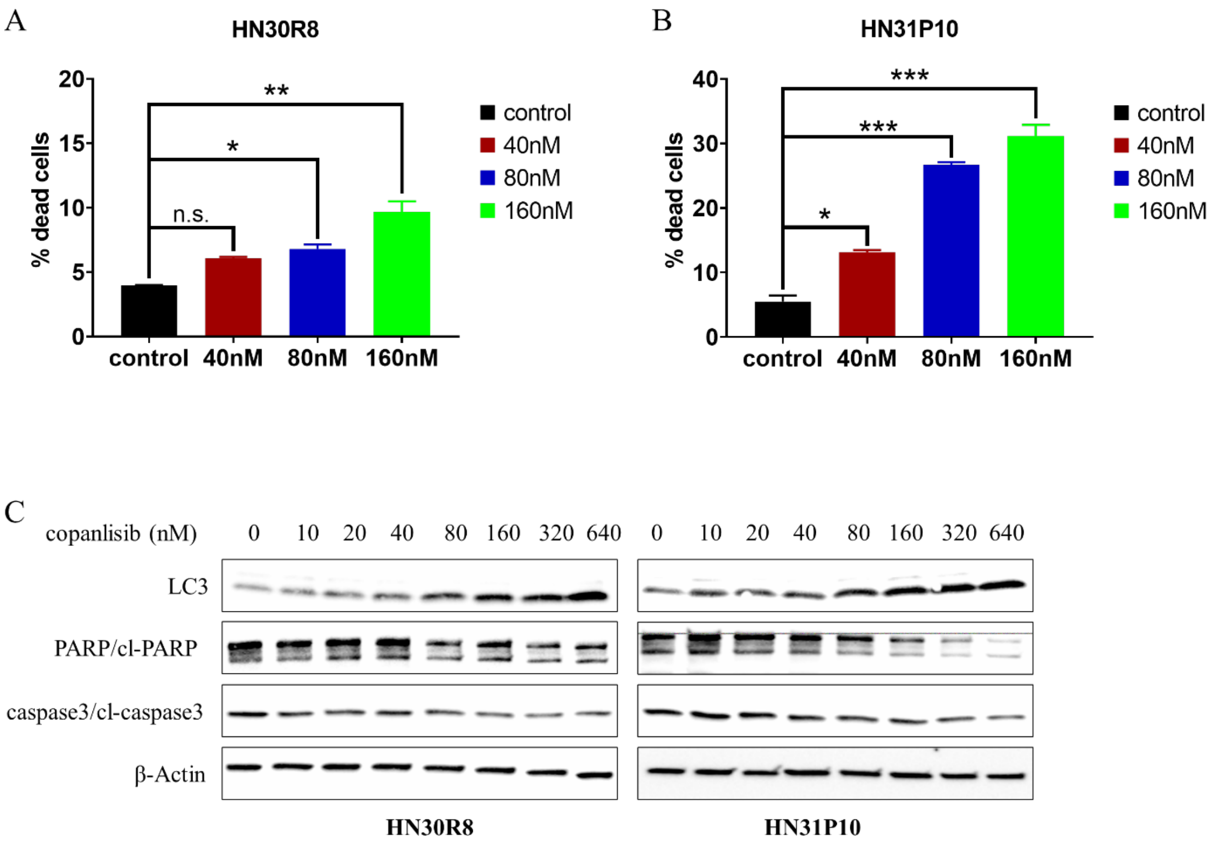


**Figure S9. Copanlisib promotes cell death in HNSCC cisplatin-resistant cell lines. (A-B)** Flow cytometry analysis shows the effect of copanlisib on cell viability in HN30R8 and HN31P10 cisplatin-resistant cell lines after 48 hours of treatment. The bar graphs display the percentage of dead cells, with the x-axis representing different copanlisib concentrations (nM) and each bar color corresponding to a specific concentration. Data are presented as mean ± standard error of the mean (SEM), with statistical significance determined by the one-way ANOVA, Dunnett's test (not significant (n.s.) $*P \leq 0.05$, $**P \leq 0.01$, $***P \leq 0.001$). **(C)** Western blot analysis of protein expression in HN30R8 and HN31P10 cells following 48-hour treatment with copanlisib. β-actin was used as a loading control to ensure equal protein loading across samples.

**Figure S10**

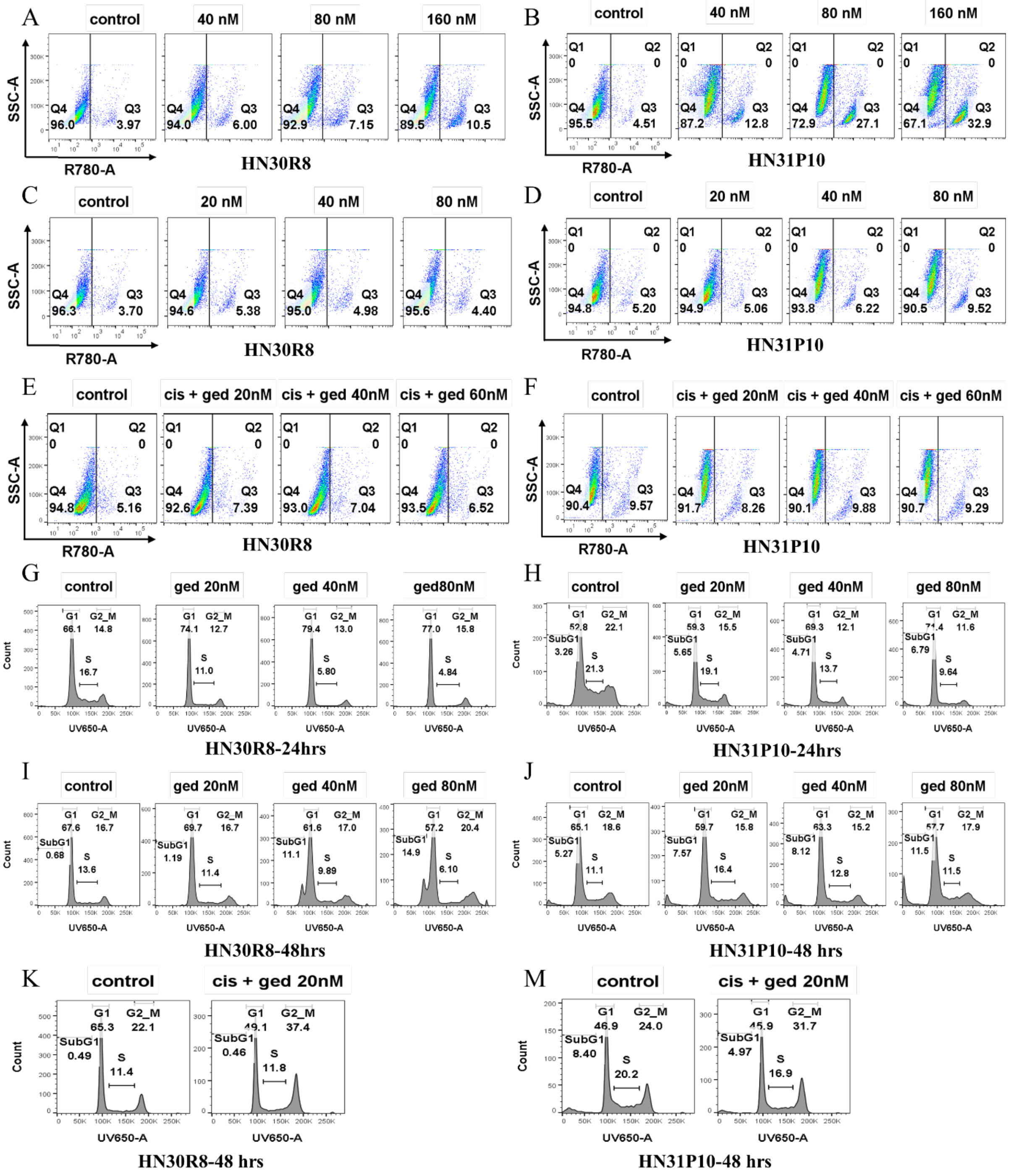


**Figure S10. Comparative analysis of PI3K inhibitors effect on cell death and cell cycle. (A-B)** Flow cytometry panels depicting cell viability of HN30R8 and HN31P10 cisplatin-resistant cell lines after 48-hour treatment with varying concentrations of copanlisib. **(C-D)** Flow cytometry panels showing cell viability for HN30R8 and HN31P10 treated with different concentrations of gedatolisib for 48 hours. **(E-F)** Panels representing the combined effect of cisplatin and gedatolisib on cell viability in HN30R8 and

HN31P10. Cells were treated with a fixed concentration of cisplatin alongside varying concentrations of gedatolisib for 48 hours. **(G-H)** Flow cytometry panels analyzing the impact of gedatolisib on the cell cycle distribution in HN30R8 and HN31P10 following 24-hour treatment with different concentrations. **(I-J)** Panels illustrate the cell cycle effects after 48-hour exposure to gedatolisib, comparing cell cycle arrest patterns in HN30R8 and HN31P10. **(K-M)** Analysis of the combined treatment of cisplatin and gedatolisib on cell cycle progression in HN30R8 and HN31P10 after 48 hours

**Figure S11**

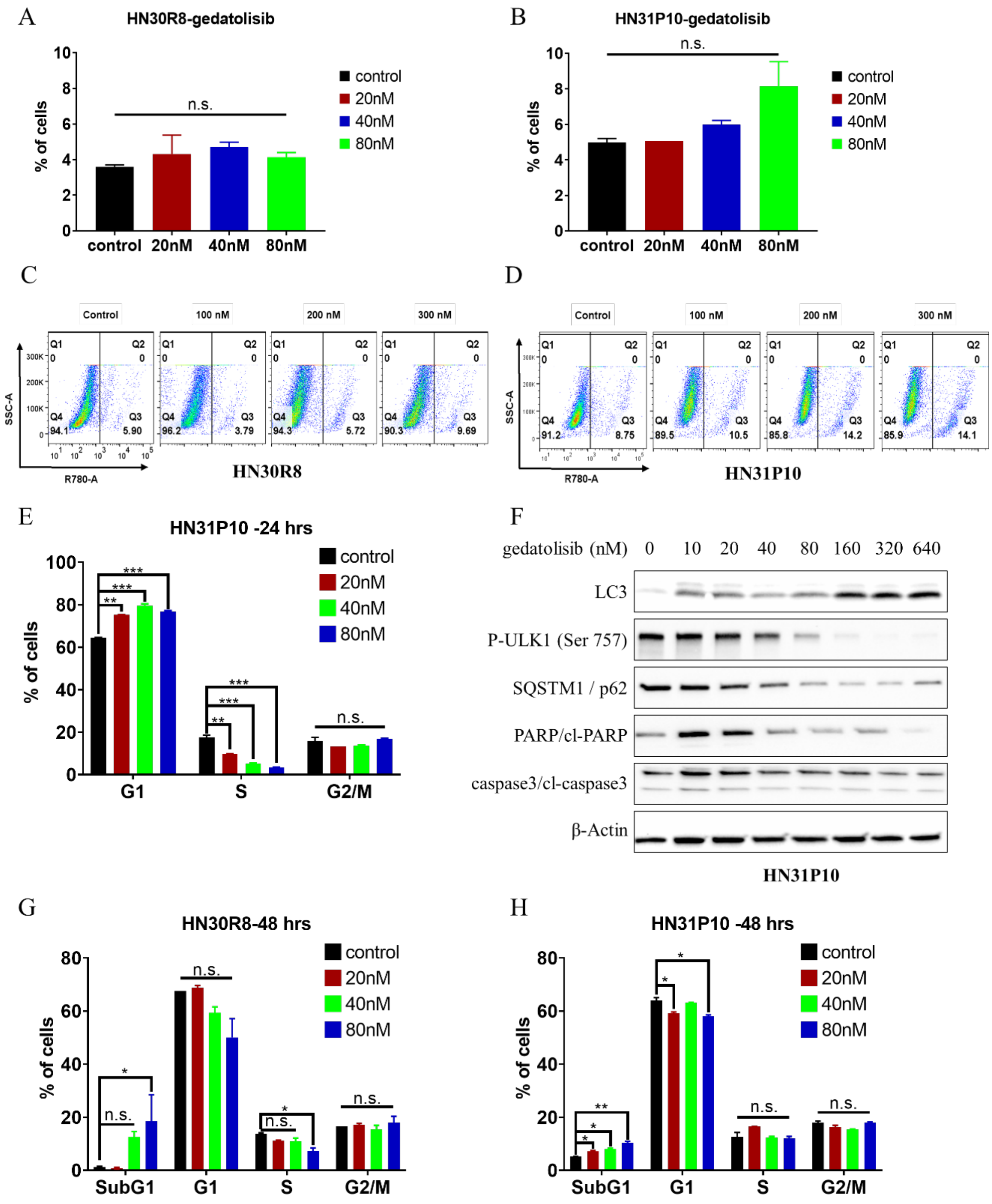


**Figure S11. Gedatolisib does not drive caspase-induced cell death. (A-B)** Flow cytometry analysis shows the effect of gedatolisib on cell viability in HN30R8 and HN31P10 cisplatin-resistant cell lines after 48 hours of treatment. The bar graphs display the percentage of dead cells, with the x-axis representing different gedatolisib concentrations (nM) and each bar color corresponding to a specific concentration. Data are presented as mean ± standard error of the mean (SEM), with statistical significance determined by the one-way ANOVA, Dunnett’s test (not significant (n.s.). (C-D) Flow cytometry panels showing cell viability for HN30R8 and HN31P10 treated with different concentrations of gedatolisib for 48 hours. **(E)** Flow cytometry cell cycle analysis for HN31P10 treated with gedatolisib in a concentration-dependent manner over 24 hours. The y-axis represents the percentage of cells in different cell cycle phases, with the x-axis indicating copanlisib concentrations (nM). Each bar color corresponds to a distinct cell cycle phase. Data are expressed as mean ± SEM, with statistical analysis conducted comparing gedatolisib treated groups to the control groups (0 nM) using an unpaired, two-tailed Student's t-test (*P ≤ 0.05, **P ≤ 0.01, ***P ≤ 0.001, ****P ≤ 0.0001). **(F)** Western blot analysis of protein expression in HN31P10 cells following 48-hour treatment with gedatolisib. β-actin was used as a loading control. **(G-H)** Flow cytometry cell cycle analysis for HN30R8 and HN31P10 treated with gedatolisib in a concentration-dependent manner over 48 hours. The y-axis represents the percentage of cells in different cell cycle phases, with the x-axis indicating gedatolisib concentrations (nM). Each bar color corresponds to a distinct cell cycle phase. Data are expressed as mean ± SEM, with statistical analysis conducted using an unpaired, two-tailed Student's t-test (*P ≤ 0.05, **P ≤ 0.01, ***P ≤ 0.001, ****P ≤ 0.0001).

#### Figure S12

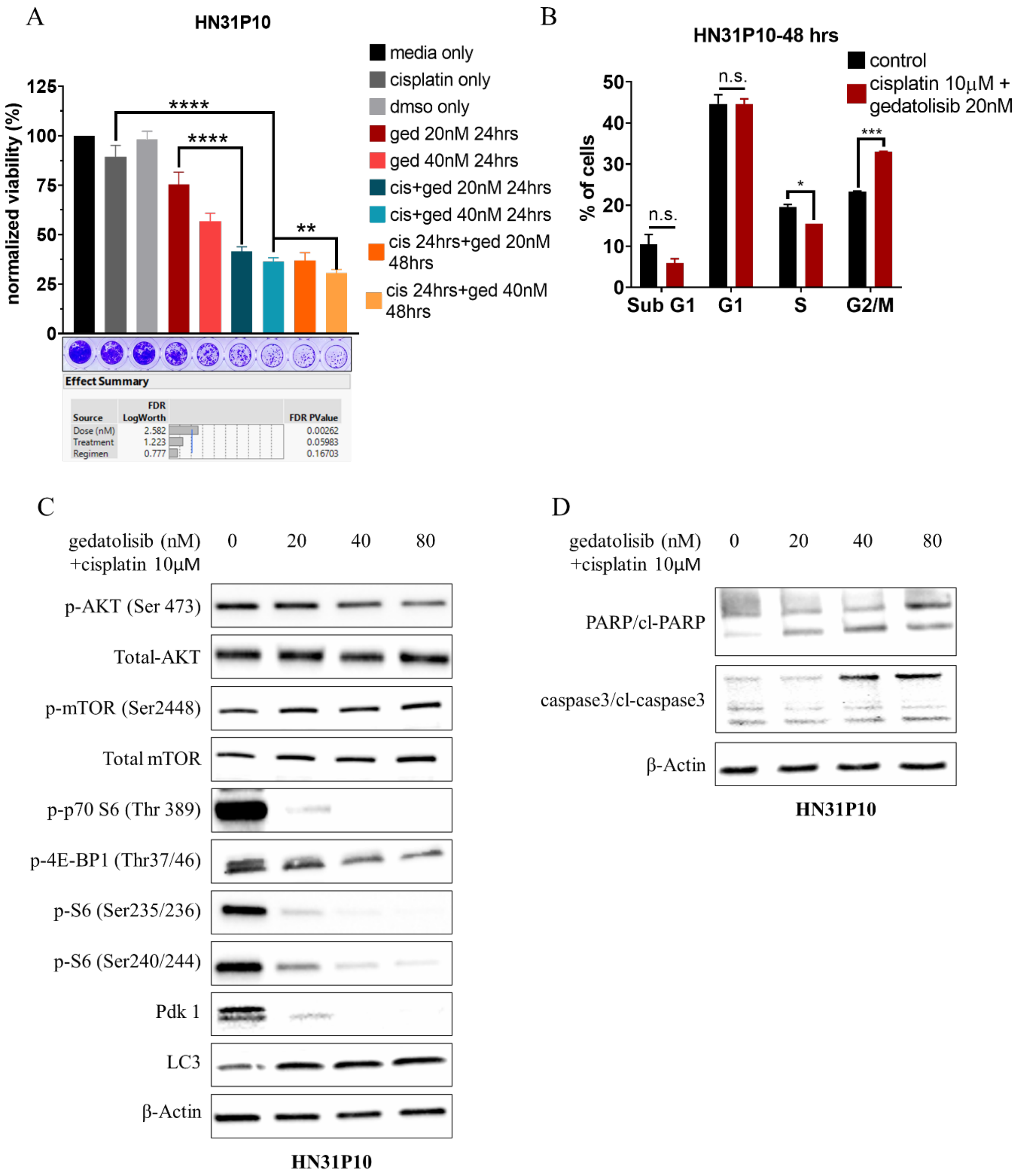


**Figure S12. Combining gedatolisib and cisplatin leads to G2/M cell cycle arrest. (A)** Activity of gedatolisib and cisplatin at varying concentrations for different timepoints in HN31P10. The bar graph represents the cell viability of the cells obtained from same samples and normalized to day 0. The y-axis shows percentage of cell viability and x-axis represents the respective samples. Data are presented as the mean

± standard error of the mean (SEM). P-values were calculated using a three-way ANOVA with Tukey's test (** P ≤ 0.01, **** P ≤ 0.0001).). **(B)** Flow cytometry cell cycle analysis bar graph showing sub G1, G1, S, and G2/M phases in HN31P10 treated with combination of gedatolisib and cisplatin for 48 hours. The y-axis represents the percentage of cells in each respective cell cycle phase, while the x-axis indicates the cell cycle phases. Black represents cisplatin alone, while red represents the combination of gedatolisib and cisplatin. Data are presented as the mean ± standard error of the mean (SEM). P-values were calculated using a one-way ANOVA with Dunnett's test (not significant (n.s.), * P ≤ 0.05, ***P ≤ 0.001). **(C-D)** Western blot panel for HN30R8 treated with combination of gedatolisib in a dose dependent manner and constant cisplatin concentration. β-actin was used as a loading control.

#### Figure S13

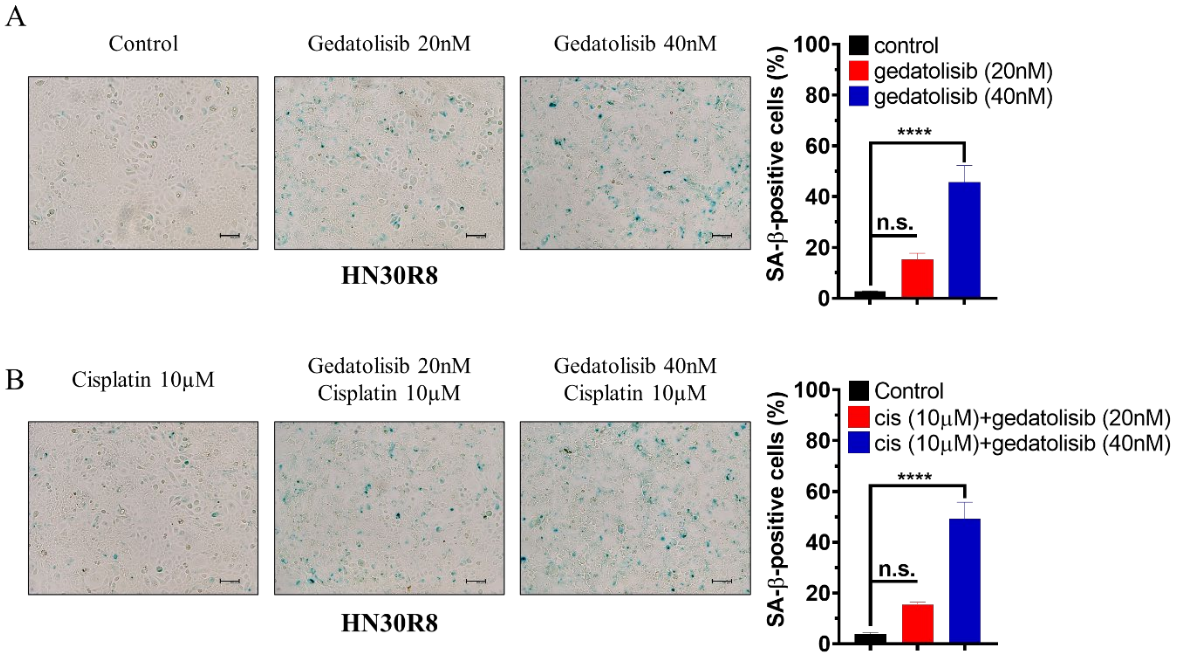


**Figure S13. Gedatolisib induced cellular senescence. (A)** HN30R8 cell line stained for β- galactosidase for control, gedatolisib treated with 20nM and 40nM, respectively. **(B)** HN30R8 cell line stained for β-galactosidase for control, gedatolisib treated with 20nM and 40nM, respectively along with 10μM cisplatin. The bar graph illustrates the percentage of positively stained senescent cells in samples treated with gedatolisib alone and in combination with cisplatin. The y-axis represents the percentage of senescence-associated positive cells relative to the total cell count, while the x-axis denotes the respective treatment groups. Data are presented as the mean ± standard error of the mean (SEM). Statistical significance was determined using a one-way ANOVA with Dunnett's test, with P-values indicated as follows: not significant (n.s.) and **** P ≤ 0.0001.

### Figure S14

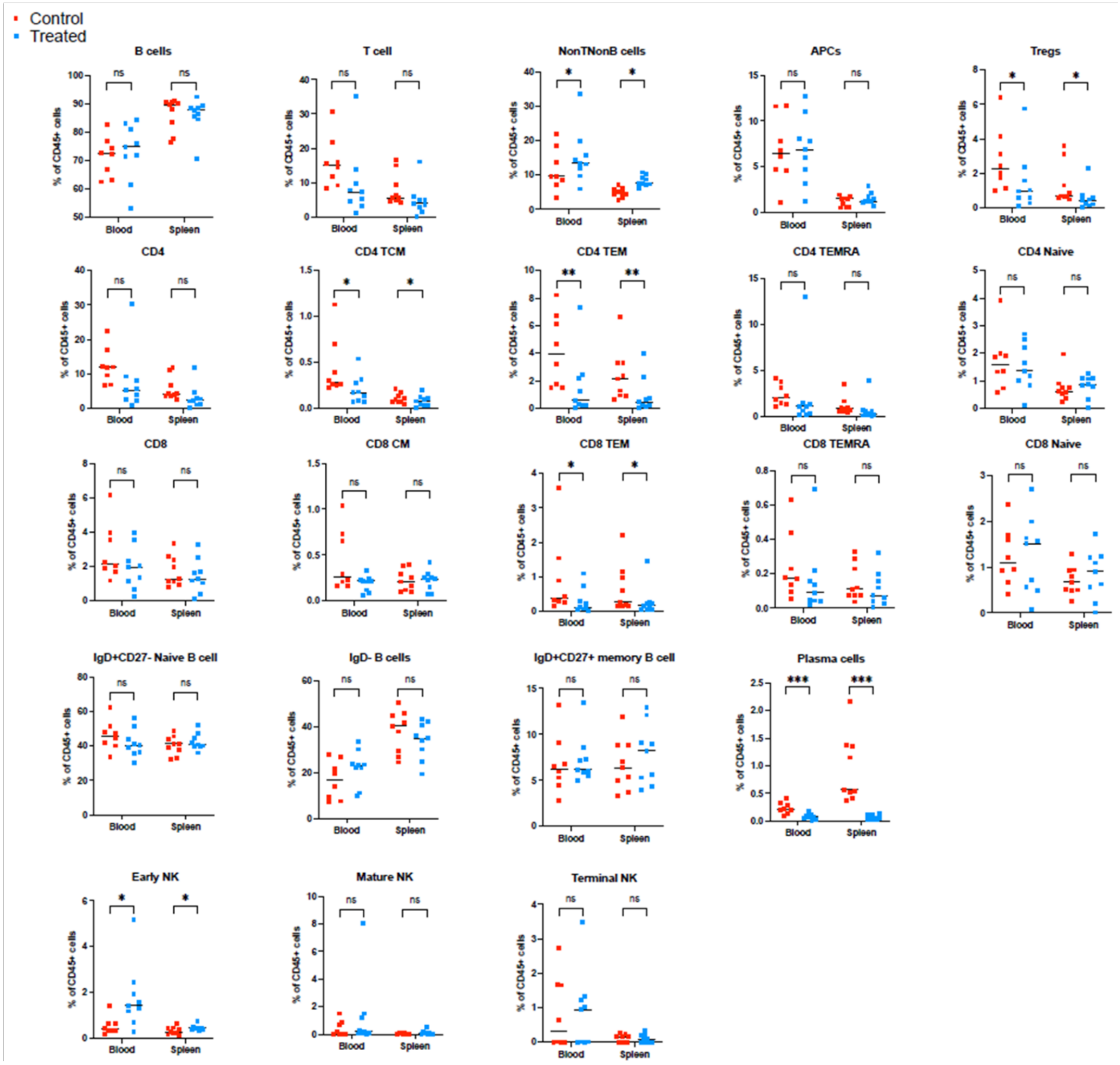


**Figure S14. Systemic effects of gedatolisib. (A)** Flow cytometry panels representing bar graph for each subclass of immunocytes in blood and spleen as a percentage of human CD45$^{+}$ cells in each tissue. The y-axis represents the frequency of immunocytes, while the x-axis indicates sample groups. Data are expressed as the median. Statistical significance was determined using a two-way ANOVA test, with significance levels denoted as follows: ns (not significant), *P ≤ 0.05, **P ≤ 0.01, and ***P ≤ 0.001.

**Figure S15**

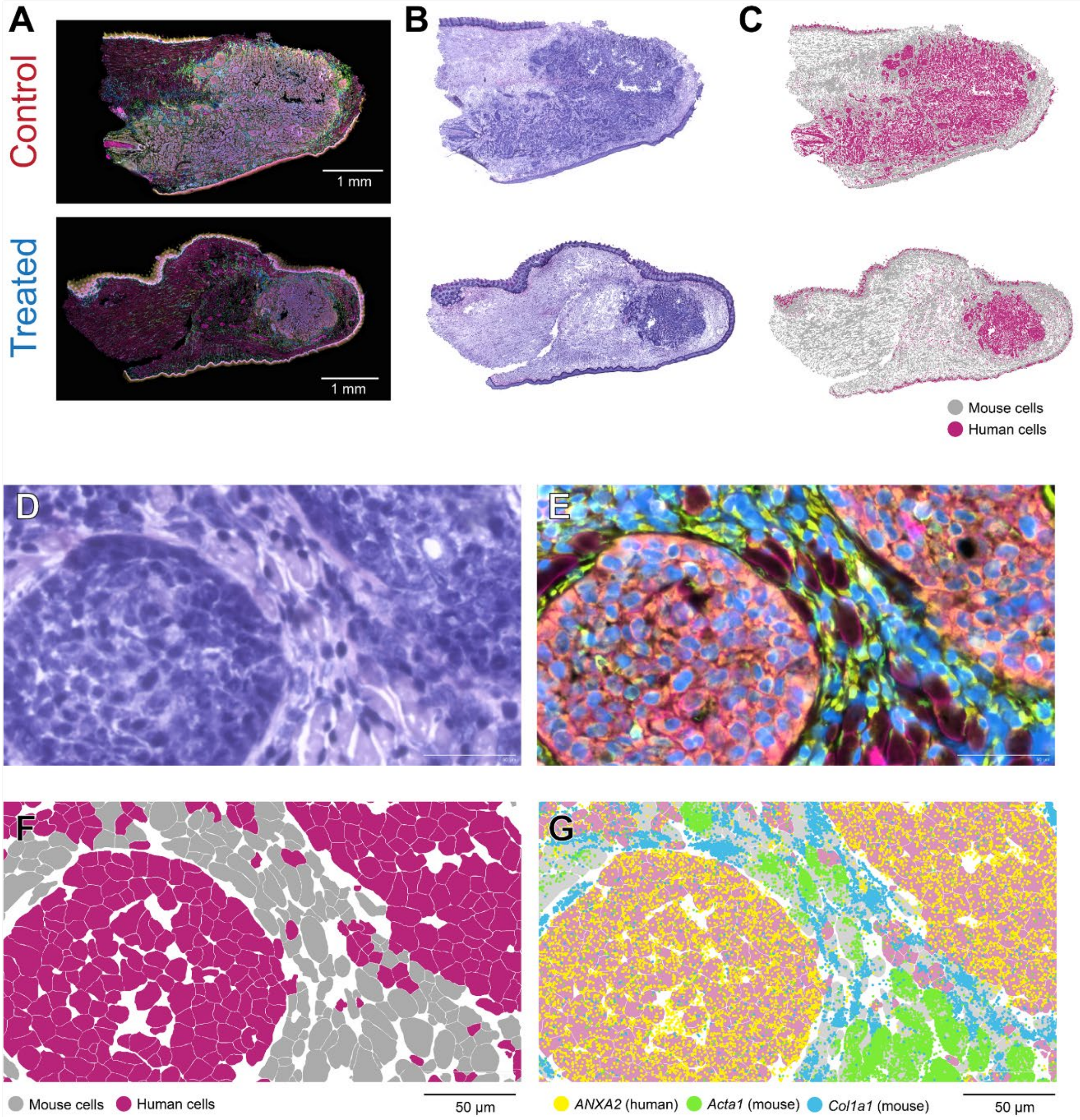


**Figure S15. Spatial differentiation of mouse and human cells. (A)** Control and treated tumors with comprehensive overlapping spatial mapping. **(B)** Conventional H&E of murine tumors in cross section. **(C)** Gene expression differentiates murine and human cells. High magnification H&E **(D)**, individual cell types **(E)**, murine vs human cells **(F)** and individual gene expression **(G)** demonstration.
